# Asparagine couples group A *Streptococcal* metabolism to virulence

**DOI:** 10.1101/2024.07.08.602371

**Authors:** Abhinay Sharma, Aparna Anand, Miriam Ravins, Nicola Horstmann, Kevin S. McIver, Emanuel Hanski

## Abstract

*Streptococcus* (GAS) is a highly adapted and human-restricted pathogen causing a wide variety of infections, some life-threatening^1^. This ability is linked to the expression of many virulence factors, whose transcription is regulated by the two-component system, CovR/S^2–5^.

Here, we show that genome transcription of GAS cultured in a chemically defined medium (CDM) is globally affected when supplemented with asparagine (Asn), including increased expression of many virulence genes. For the first time, we report that GAS solely depends on asparagine synthetase (AsnA) for Asn synthesis, on the ABC transporter (GlnPQ) to import Asn, and on the asparaginase (AsnB) to maintain a precisely balanced intracellular Asn concentration. Furthermore, we show that mutants defective in either *asn*A, *gln*P, or *asn*B express significantly lower levels of virulence factors in CDM and are severely attenuated in the sublethal murine model of human GAS soft-tissue infection.

We further show that the synthesis and import of Asn in GAS are ATP-dependent and negatively regulated by intracellular Asn. Thus, Asn availability controls the intracellular ATP level. When ATP becomes limiting, CovR phosphorylation decreases. This augments GAS growth rate, virulence production, metabolism, and *vice versa* when the ATP level increases. Furthermore, excess Asn accumulates inside GAS in AsnB mutant, destroying the balance between Asn and ATP. We discuss the high similarity between these mechanistic principles of the Asn-mediated control of GAS virulence and metabolism to the Asn-mediated control of tumor growth^6^, indicating evolutionary significance.

## Introduction

*Streptococcus pyogenes* - Group A streptococcus (GAS) is an extracellular strict human pathogen recognized among the top ten causes of mortality from infections^7^. It may exist as a benign colonizer^8^. However, it also causes many human manifestations, including life-threatening infections such as bacteremia, necrotizing fasciitis (NF), and streptococcal toxic shock syndrome STSS^8, 9^. In addition, GAS may trigger a lethal autoimmune sequela^10, 11^. In 2022-2024, a surge in invasive GAS diseases in Europe, the USA, and Japan was observed^12–15^.

Bacterial pathogens respond to specific nutritional cues within host microenvironments, and growth within these microenvironments requires specific metabolic pathways. Studies along these lines peaked in recent years, and novel traits have been uncovered mostly for intracellular pathogens or pathogens occupying specific host niches^16^. However, we know little about extracellular pathogens, such as GAS, that co-exist with their host or cause many diseases and pass through various host environments during the infectious process^8^. We have shown that acquiring Asn from the host by GAS during infection is a critical property of GAS virulence^17^. GAS delivers the toxins streptolysin S (SLS) and streptolysin O (SLO) into infected cells. These toxins cause endoplasmic reticulum (ER) stress, activating the PERK-eIF2α-ATF4 branch of the unfolded protein response (UPR)^18^. The activation of the PERK-eIF2α-ATF4 upregulates the host asparagine synthetase (ASNS) transcription^19^; consequently, the Asn level is augmented in infected host cells^17, 18^. GAS utilizes the generated Asn to increase its growth rate and degree of virulence^17^. By inhibiting the PERK-eIF2α-ATF4 pathway using specific inhibitors, we protected mice against invasive GAS diseases in murine models of soft tissue GAS infection^18^.

The best-characterized primary regulator of GAS virulence is the two-component system (TCS) termed control of virulence (CovR/S) or capsule synthesis regulator (CsrR/S)^3, 20^. CovR represses virulence factors transcription when phosphorylated^3^. CovS acts as a kinase and phosphatase to modulate CovR phosphorylation levels; the human cathelicidin host-defense peptide (LL-37) binds to CovS and increases its phosphatase activity^21, 22^.

Whereas the requirement for host Asn in invasive GAS infection is well documented, how Asn regulates GAS virulence has remained elusive. This study addressed this knowledge gap. Here, we show, for the first time, that Asn synthetase (AsnA), the Asn importer (GlnPQ), and asparaginase (AsnB) control intracellular Asn concentration. AsnA and GlnPQ activities are ATP-dependent processes that consume ATP while maintaining the appropriate Asn concentration. AsnB effectively degrades Asn, which controls AsnA and GlnpQ transcription through negative feedback inhibition. The resulting limited ATP availability decreases CovR phosphorylation, thereby increasing GAS virulence growth rate and metabolism. Mutants deficient in AsnA, GlnPQ, and AsnB activities are attenuated in the sub-lethal murine model of human GAS soft-tissue infection, demonstrating that this exquisite mode of regulation occurs *in vivo* during infection. Here, we discuss the high similarity between the mechanistic principles of the Asn-mediated control of GAS virulence and metabolism to the Asn-mediated control of tumor growth.

## Results

### Detection of Asn-mediated regulatory circuits by RNA-seq

This study mainly used the M1T1 S119 invasive GAS strain isolated from a patient with NF and STSS^23, 24^. To decipher the effect of Asn on GAS gene transcription, we performed transcriptome sequencing (RNA-seq) experiments in the chemically defined medium (CDM) designed by van de Rijn and Kessler, which initially did not contain Asn^25^ because GAS is not auxotrophic to Asn. Supplementing the CDM with Asn at increasing concentrations of 0.5 to 10 µg/ml enhanced the S119 growth rate, reaching the fastest growth at 10 µg/ml (Fig. 1a). Nevertheless, after prolonged growth (overnight), the cultures with and without Asn reached similar optical density (Fig. 1a). The RNA-seq experiment compared RNAs of S119 grown to an OD_600_ = 0.7 in CDM in the presence and absence of 10 µg/ml Asn. The heatmap of the genes having 2-fold higher or -lower transcript abundance shows that they belong to several functional categories: Cell communications, Cell division/replication proteins, Hypothetical proteins, Kinases, Metabolic enzymes, Phage-associated proteins, Regulatory proteins, Stress related proteins, Transferases, Transporters/ABC transporters/solute binding proteins, and Virulence factors (Fig. 1b). Thus extracellular Asn present at low concentration of 75 µM (10 µg/ml) exerts a global effect; 27 % (n = 503) of genes were affected; out of these, 13.97% (n = 257) were upregulated, and 13.37% (n = 246) were down-regulated in the presence of Asn. Among the most upregulated genes were SHP2 and SHP3, encoding autoinducing peptide pheromones of the quorum sensing Rgg2/3 pathways leading to increased biofilm formation and resistance to the bactericidal effects of the host factor lysozyme^26^. In addition, glucose-6 phosphate isomerase (a highly conserved glycolytic enzyme), glycosyltransferase (involved in cell wall biogenesis), and the eukaryotic-type serine/threonine kinase (involved in cell cycle regulation)^27^ were also found highly upregulated (Extended Data Figs. 1a, b). The genes most strongly downregulated encoded mainly unknown proteins (Extended Data Figs. 1a, b).

**Fig. 1.**
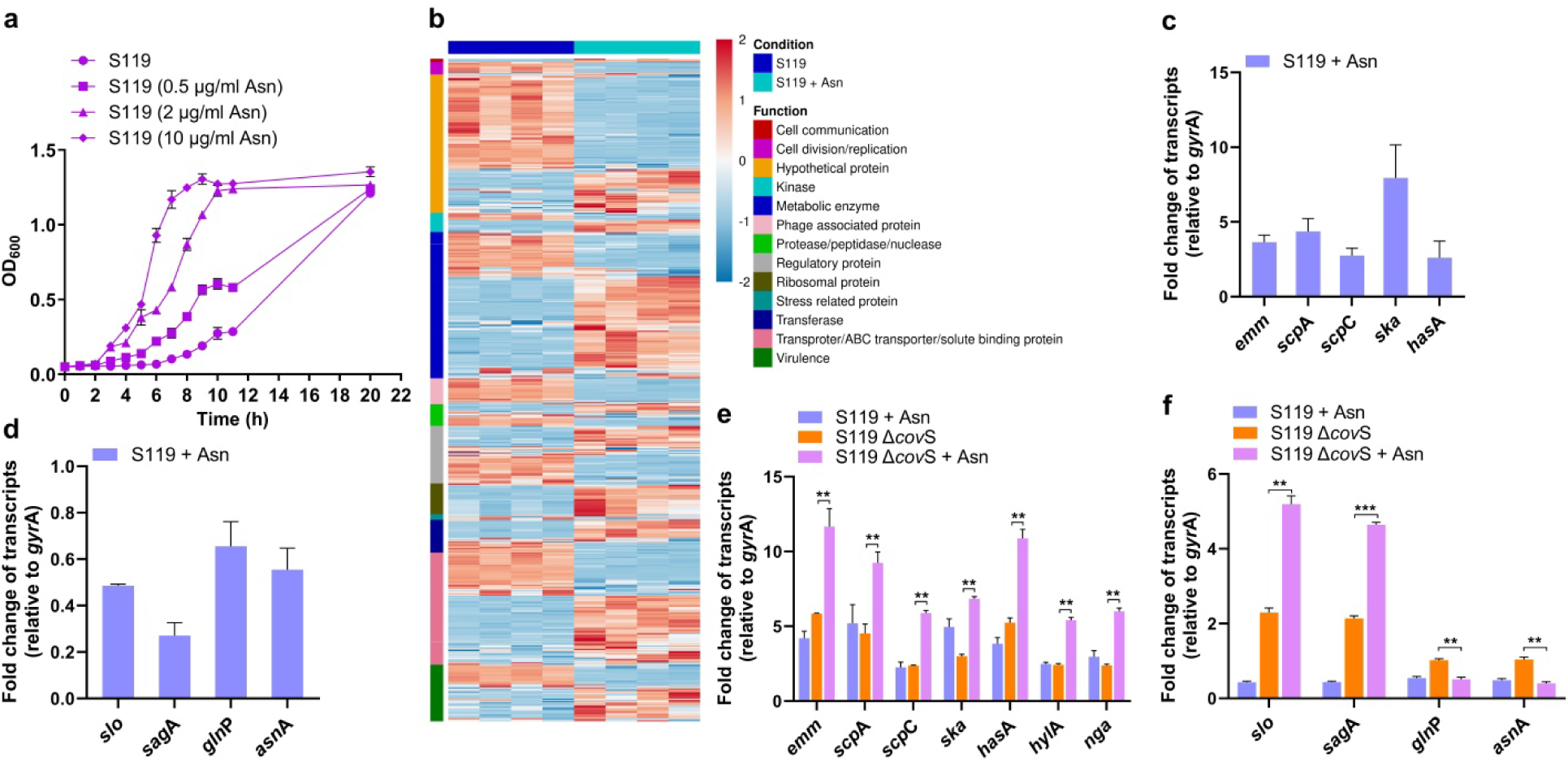
Asn affects GAS growth and transcription. **a,** The growth of the GAS strain S119 was determined in CDM in the absence or presence of Asn (0.5, 2, 10 µg ml^-1^). The values shown represent the means ± S.D. *n=2*. **b,** The heatmap shows differential gene expression patterns based on RNA-seq data. Data illustrates the global differential expression of genes belonging to the indicated different functional categories. *n=4*. **c,d,** Quantitative real-time PCR (qRT-PCR) validations of mRNA-seq data were performed. Upregulated (*emm*, *scp*A, *scp*C, *ska,* and *has*A) (**c**), and down-regulated genes (*slo*, *sag*A, *gln*P, and *asn*A) (**d**), are presented. The determinations were performed on the two RNA samples used for the RNAseq experiment. *n=2*. **e,f,** qRT-PCR determinations of Asn effect on the transcription of selected genes (*emm*, *scp*A, *scp*C, *ska*, *has*A, *hyl*A, *nga*) Set 1 (**e**), (*slo,* and *sag*A, *asn*A*, gln*P) Set 2 (**f**), in S119 and its derived Δ*cov*S mutant in CDM without or with Asn (10 µg ml^-1^). *n=2*. In all qRT-PCR data, transcript abundance for each gene was normalized to that of *gyr*A in each sample, and fold change was calculated in comparison with the normalized transcript abundance of the S119 grown without Asn (**c-f**). The values shown represent the means ± S.D. Statistical analysis was performed using an unpaired two-tailed t-test; **P < 0.01 ***P<0.001 (**c-f**).

To confirm the results concerning virulence factors repressed by phosphorylated CovR/S^21, 28, 29^, such as *emm*, *scp*A, *scp*C (a.k.a.s*pyCEP*), and *has*A, we used quantitative reverse transcriptase-polymerase chain reaction (qRT-PCR) determinations under the same conditions. We found that the CovR/S regulon genes were significantly upregulated in the presence of Asn, suggesting that Asn might decrease CovR/S-mediated repression of virulence factors transcription (Fig. 1c). In addition, AsnA transcription was downregulated, suggesting that Asn acts as an allosteric feedback inhibitor of AsnA^30^ (Fig. 1d). Nevertheless, the transcription of SLO and SLS, both known to be repressed by CovR/S^31^, were repressed under these conditions (Fig, 1d). We assume they might also be controlled by the same feedback inhibitory circuit regulating *asnA* because they generate ER stress and UPR response in the infected host cells, which produce Asn that the bacterium utilizes^17^. Thus, when Asn is present in the medium, this circuit is inhibited to maintain the desired range of intracellular Asn concentrations.

To test how mutations in CovR/S would affect the Asn-mediated regulation, we constructed a mutant deficient in CovS (Δ*cov*S) and performed the qRT-PCR determinations again (Fig. 1e, f). As shown previously^32^, the transcription of all tested CovR/S-controlled genes was upregulated in the mutant compared to WT. However, Asn could still upregulate most of the genes (Fig. 1e, f), and the gene encoding AsnA maintained the negative feedback inhibition (Fig.1f). However, when both covR and covS genes were inactivated, the regulated genes were no longer upregulated by Asn. In contrast, the negative regulation of AsnA by Asn was still maintained (Extended Data Fig. 1c, d). These results suggest that CovR is necessary for Asn-mediated upregulation of virulence genes.

The ATP-binding cassette transporter GlnPQ is an essential glutamine and glutamic acid uptake system and transports Asn in Gram-positive bacteria^33, 34^. Its transcription is strongly upregulated in the culture medium of the M1T1 strain MG5005 containing the Gram-negative asparaginase, Kidrolase that depletes Asn from the medium^17^, and was also upregulated in the S119 strain in the absence of Asn, albeit below the 2-fold higher threshold. The qRT-PCR results show that *gln*P transcript was significantly more abundant when S119 was cultured in the absence of Asn than in its presence (Fig. 1d), suggesting that GlnPQ could participate in the regulatory circuit of SLO, SLS, and AsnA (Fig. 1d) and thus be responsible for Asn transport.

### Control of intracellular Asn

#### I. Role of AsnA

To test the role of asparagine synthetase (AsnA) in GAS growth and gene transcription, we constructed a deletion mutant in *asn*A (Δ*asn*A). We found that Δ*asn*A is Asn-auxotrophic when cultured in CDM (Fig. 2a). Its maximum growth rate could be restored by supplementing the CDM with Asn or Ala-Asn dipeptide (Fig. 2a), suggesting that Asn cannot be formed by Δ*asn*A when cultured in CDM, but extracellular Asn or Ala-Asn supports growth when brought in *via* Asn importer or likely through the dipeptide permease^35^. In addition, the increase in the growth rate of the AsnA-deficient mutant was restored by genetic complementation with a WT *asn*A gene expressed from a plasmid in CDM without Asn (Extended Data Fig. 2a). A plasmid expressing a mutated AsnA containing a single amino acid mutation that rendered it catalytically inactive (replacement of arginine with lysine at amino acid 100, Arg100Lys) did not complement the growth without Asn (Extended Data Fig. 2a).

**Fig. 2.**
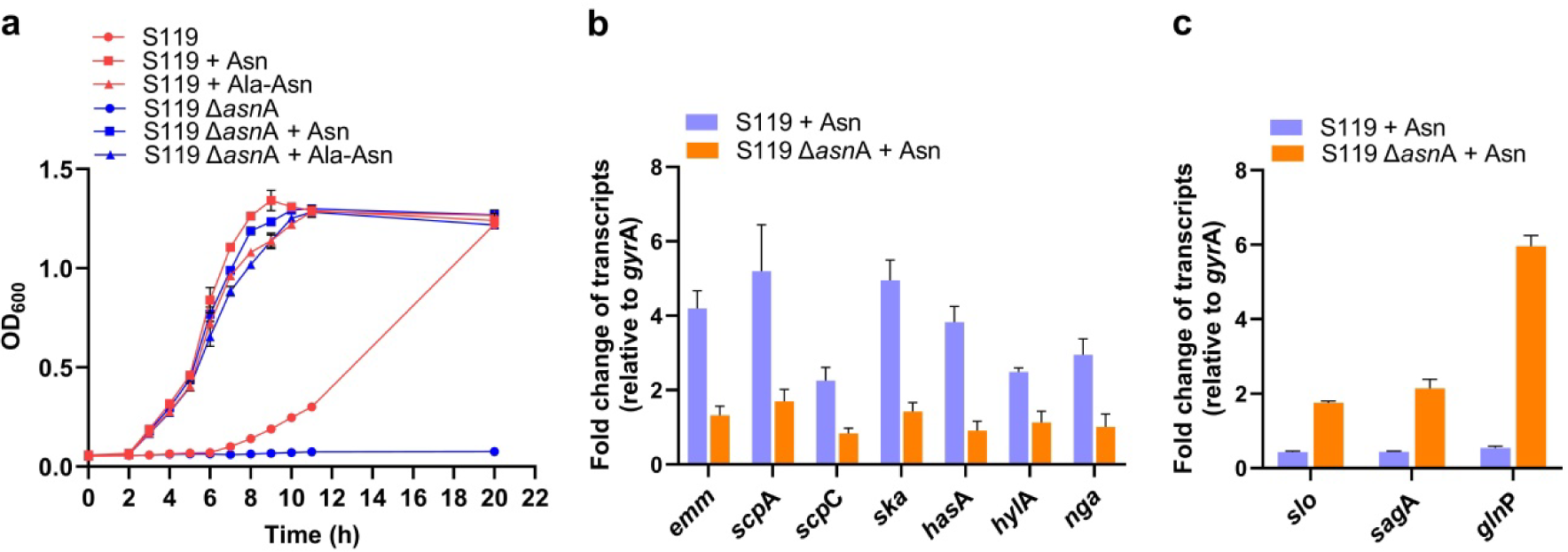
*asn*A is essential for Asn-mediated gene regulation. **a,** The strains S119 and S119Δ*asn*A were grown in CDM in the absence or presence of Asn (10 µg ml^-1^) or dipeptide (Ala-Asn) (100 µg ml^-1^) and OD_600_ was determined at indicated time intervals. *n=2*. **b,c,** qRT-PCR determinations of Set 1 (**b**), and Set 2 (**c**) of S119 and S119 Δ*asn*A genes were performed as in Fig. 1. *n=2*. (**b,c)**. The data shown represent the means ± S.D. Statistical analysis was performed using an unpaired two-tailed t-test; *P <0.05 **P < 0.01 (**b,c)**.

Next, we found that the Δ*asn*A mutant lost the Asn-mediated upregulation of GAS virulence factors using qRT-PCR (Fig. 2b). However, it was restored by complementing the mutant with a plasmid expressing the WT gene but not with an Arg100Lys catalytically inactive allele, suggesting the formation of Asn is essential for the AsnA-mediated regulation (Extended Data Fig. 2b). Furthermore, the negative feedback regulation controlling the expression of the genes encoding SLO and SLS was lost in the Δ*asn*A mutant and restored by complementation with the WT gene and not by the catalytically inactive gene (Extended Data Fig. 2c). Interestingly, in the Δ*asn*A mutant, the transcription of *gln*P was increased by about 6-fold (Fig. 2c), likely to compensate for the loss of Asn production, suggesting that *gln*PQ and *asn*A regulation are tightly controlled. In addition, we demonstrated that a deletion of *asn*A in the M1T1 strain 854 exerted similar effects on its growth in CDM and gene regulation (Extended Data Figs. 2d-f). The role of RocA, known to participate in CovR regulation, was ruled out in these processes (Extended Data Figs. 2g-i).

In summary, catalytically active AsnA is necessary for Asn-mediated gene regulation but not for growth in the CDM supplemented with Asn or di-peptide-containing Asn.

#### II. Role of GlnPQ

To test the role of the ATP-binding cassette transporter GlnPQ on GAS growth and gene transcription, we constructed an insertion-inactivation mutant (*gln*P^-^) and followed its growth pattern in un-supplemented CDM or CDM supplemented with Asn. The *gln*P^-^ and WT strain S119 grew in the absence of Asn at similar slow rates, but the *gln*P^-^ mutant failed to increase its growth rate in CDM supplemented with 10 or even 100 µg/ml of Asn (Fig. 3a). Genetic complementation with WT *gln*P gene expressed from a plasmid restored the enhanced growth phenotype of the mutant in the presence of Asn (Fig. 3a). These experiments suggested that GlnPQ significantly contributes to Asn transport in GAS.

**Fig. 3.**
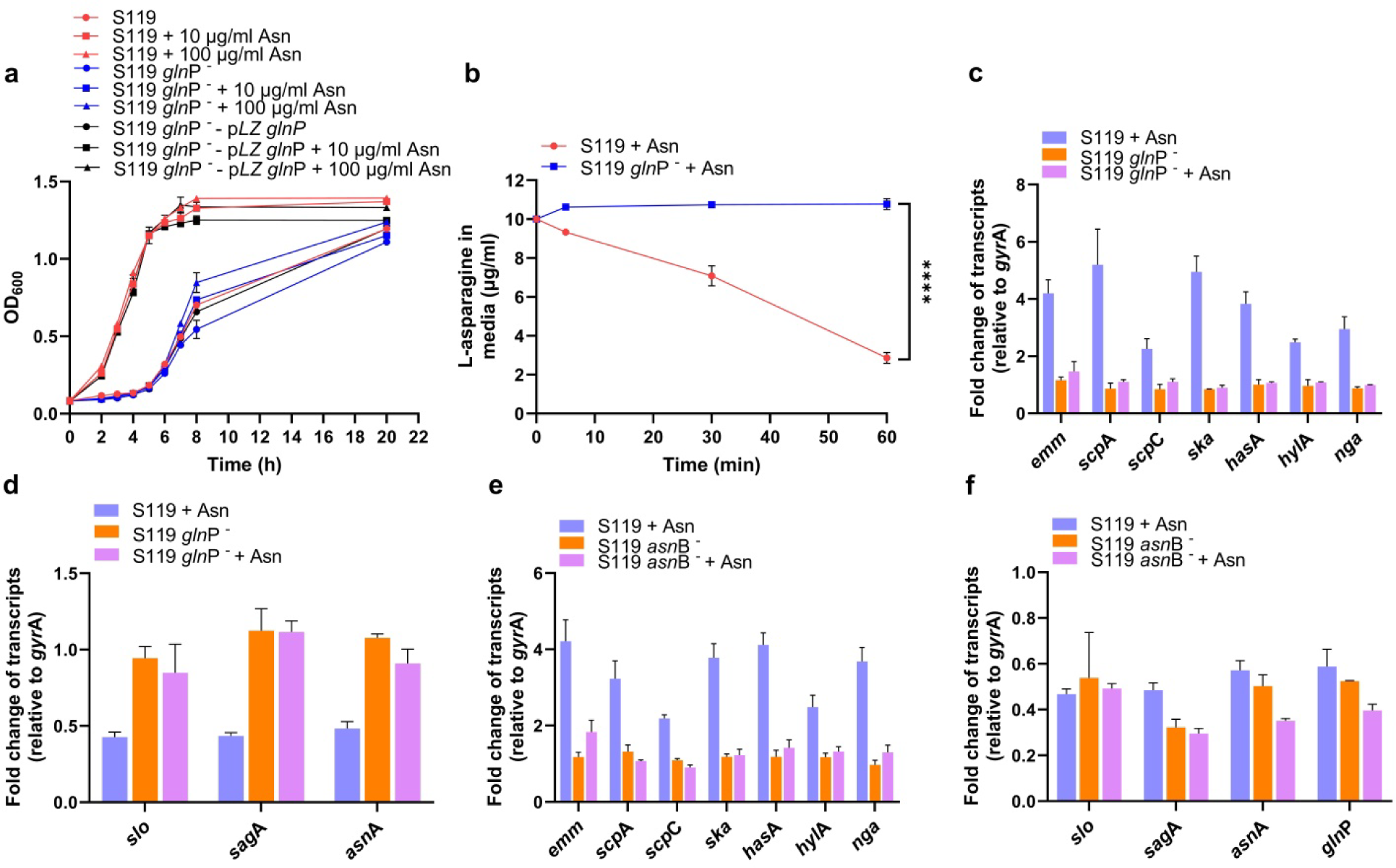
*gln*P and *asn*B are essential for Asn-mediated gene regulation. **a,** The growth of S119, S119 *gln*P^-^ and S119 *gln*P^-^-p*LZgln*P in CDM in the absence or presence of Asn (10 and 100 µg ml^-1^) was determined at indicated time points. *n=2*. **b,** S119 or its *gln*P^-^-derived mutant GAS was cultured in CDM without Asn to OD_600_ = 0.2, and then Asn was added. The Asn concentration in the medium was determined by LC-MS at 5, 30, and 60 min after Asn addition. *n=5*. Statistical analysis was performed using a Two-Way ANOVA test ****P < 0.0001. **c,d,e,f,** qRT-PCR determinations were performed on Set 1 (**c,e**) and Set 2 of genes (**d,f**) comparing S119 and S119 *gln*P*^-^* (**c,d**) and S119 and S119 *asn*B*^-^* (**e,f**) grown in CDM without or with Asn. *n=2*. In all qRT-PCR data, transcript abundance for each gene was normalized to that of the GAS S119 strain without Asn (**c-f**). The data shown represent the means ± S.D.

To provide direct evidence that GlnPQ is responsible for the uptake of Asn in GAS, we grew S119 WT or *gln*P^-^ to OD_600_ = 0.2 in CDM without Asn. Then, Asn (10 µg/ml) was added or not, and we followed the growth of the two strains. As expected from Fig. 3a, adding Asn enhanced the growth rate of S119, but not of *gln*P^-^, which maintained a relatively slow growth rate despite the presence of Asn (Extended Data Fig. 3a). To follow the uptake of Asn, we quantified the Asn concentration in the media of the two strains by liquid chromatography-mass spectrometry analysis (LC-MS)^36^. We found that Asn was taken up from the media by S119 within 60 min. In contrast, the Asn concentration in the media for the *gln*P^-^ did not change at all and was equal to that found in fresh media (Fig. 3b). This experiment demonstrates that GlnPQ is directly responsible for Asn uptake by GAS.

To assess the contribution of the deficiency in GlnPQ activity on gene regulation, we repeated the qRT-PCR determination described above. We found that the upregulation of virulence factors in the presence of Asn was lost entirely in the *gln*P^-^ mutant (Fig. 3c). The negative feedback regulation for the expression of SLO and SLS was also lost (Fig. 3d). Interestingly, the *asn*A expression in CDM in the absence and presence of Asn was similar and about 2-fold higher than that of the WT strain in the presence of Asn (Fig. 3d), presumably to compensate for the deficiency in the uptake of Asn from the medium, and supporting the notion that the expression of *asn*A and *gln*PQ is tightly regulated (Fig. 2c). Finally, expressing WT *gln*P from a plasmid restored the WT virulence gene regulation (Extended Data Fig. 3b), *slo*, *sag* (SLS), *asn*A, and *gln*PQ genes (Extended Data Fig. 3c) confirming the role of *gln*PQ in the regulation.

#### III. Role of AsnB

Although the transcription of GAS asparaginase (AsnB) was not affected by Asn’s absence or presence in CDM or by deficiency of AsnA or GlnPQ activity (Extended Data Fig. 3d), we decided to construct an *asn*B^-^ mutant and examine its effect on GAS growth and gene regulation. The main reason was that AsnB belongs to the type II asparaginase family, having kcat and K_M_ ranging around 12–60s^-1^ and 10–20 μM, respectively^37^. Therefore, AsnB should leverage intracellular Asn concentrations due to its high efficiency. To confirm the expected *asn*B^-^ phenotype, we measured L-asparaginase activity in bacterial-cell suspensions^38^. *asn*B^-^ lost its asparaginase activity, which was restored when complemented by the WT gene expressed from a plasmid (Extended Data Fig. 3e). The deficiency in asparaginase activity increased the GAS growth rate in CDM in the absence of Asn (Extended Data Fig. 3f). However, the transcription of virulence factors in *asn*B^-^ was significantly reduced and became independent of Asn (Fig. 3e). Similarly, the transcription of the genes encoding SLO, SLS, AsnA, and GlnPQ was reduced compared to the WT S119 (Fig. 3f). Furthermore, upon genetic complementation with a WT gene expressed from a plasmid, the Asn-mediated regulation of virulence factors and the genes encoding SLO, SLS, and AsnA was regained (Extended Data Fig. 3g, h). These findings suggested that Asn-mediated transcription regulation occurs at a precise Asn intracellular concentration range, which is maintained by the interplay between the Asn intracellular concentration and its effect on the transcription of *asn*A and *gln*PQ and the impact of the transcription of the latter genes on each other (Extended Data Fig. 3i).

### Asn increases virulence factor expression by reducing CovR phosphorylation

To corroborate that an Asn-mediated increase in the transcription of virulence factors also increases the expression of the related proteins, we tested the activities of ScpC (a.k.a SpyCEP) and ScpA encoding the CXC-chemokine serine protease and C5a peptidase, respectively^39–41^. ScpC cleaves interleukin-8 (IL-8), and ScpA cleaves the complement component 5a (C5a); the two cleavage processes can be visualized on SDS-PAGE^40, 41^. In addition, it has been shown that LL-37 binds to CovS and stimulates its phosphatase activity, thereby diminishing CovR phosphorylation and consequently increasing ScpC and ScpA expression ^21, 28^. Therefore, we first tested if growing S119 in CDM in the absence or presence of Asn and or LL-37 would affect the activities of ScpC and ScpA. We collected the media at OD_600_ = O.7 and tested for cleavage of IL-8 or C5a peptides. We found a low ScpC activity for S119 grown in the CDM only. In the presence of either Asn or LL-37, some cleavage of IL-8 was visualized. However, a considerable cleavage was apparent when S119 was grown in CDM containing both Asn and LL-37 (Fig. 4a). We repeated this experiment using the S119Δ*cov*S mutant and found some cleavage when cultured in CDM alone or CDM containing LL-37. In sharp contrast, we detected a high amount of cleavage of IL-8 when S119Δ*cov*S was cultured in CDM containing either Asn or Asn and LL-37 (Fig. 4b). Similar results were obtained for S119 and its derived Δ*cov*S mutant when tested for cleavage of C5a (Extended Data Fig. 4a, b), indicating that Asn-mediated increase of virulence factors transcription also upregulates expression activity that does not require an intact CovS.

**Fig. 4.**
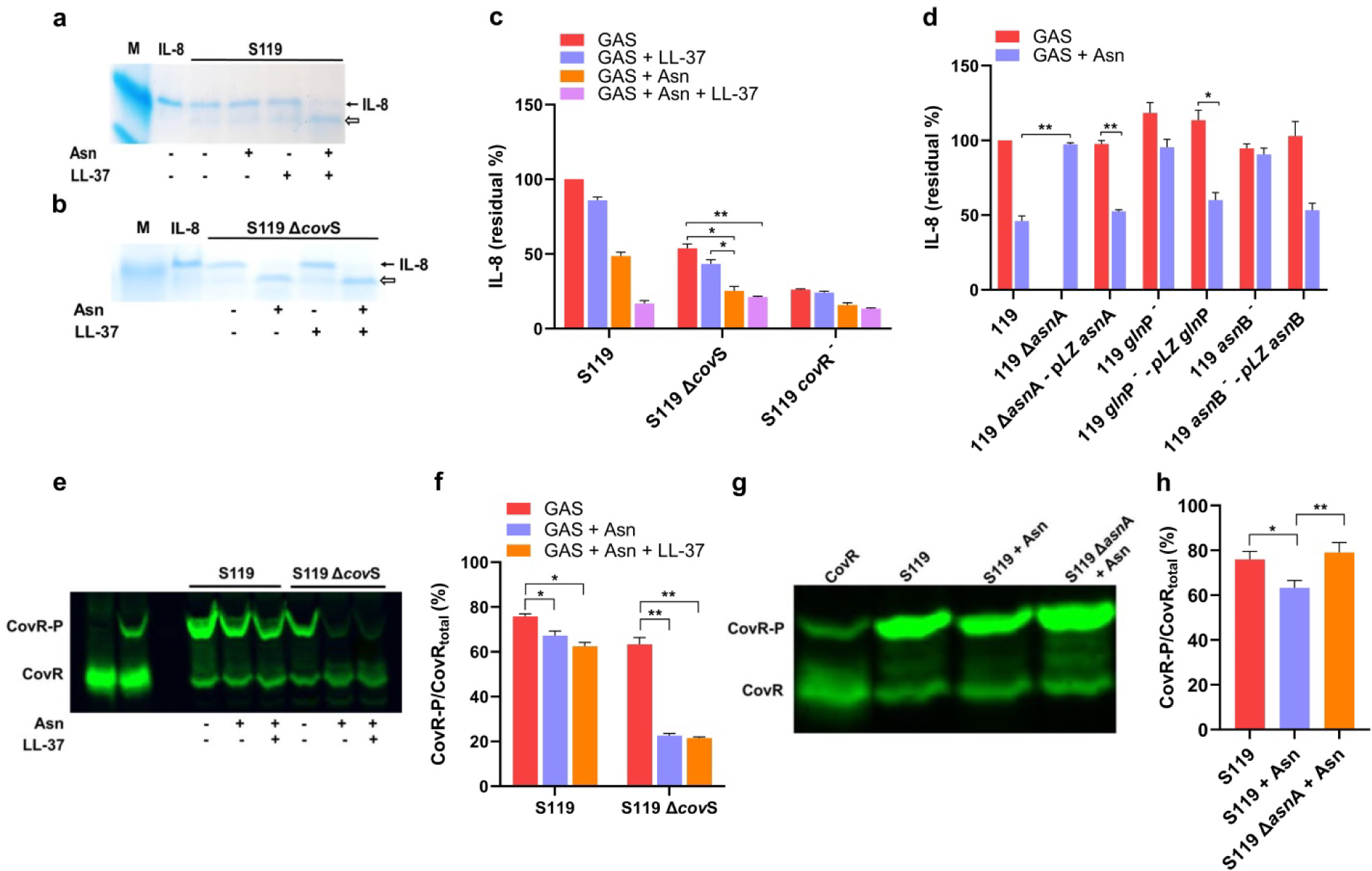
Asn reduces the phosphorylation CovR. a,b, ScpC activity in culture media. Culture media of S119 (**a**), or its Δ*cov*S-derived mutant (**b**), were collected after growth in the absence or presence of Asn or/and LL-37 and then subjected to ScpC-mediated cleavage of recombinant human IL-8 followed by SDS-PAGE on Tris-tricine gels. The gels were visualized using Coomassie blue staining. The data are representative of 2 independent experiments. *n=2*. **c,d,** The determinations of IL-8 residual content in control (100%) (supernatant of S119 in the absence of Asn) and the supernatants of the indicated strains cultured without or with Asn or/and LL-37 were conducted by ELISA. *n=2*. The data shown represent the means ± S.D. **e,g, Asn reduced CovR phosphorylation.** The indicated strains were grown in CDM without or with Asn or/and LL-37. Cell lysates (**e,g**) were separated by Phos-Tag SDS-PAGE, with unphosphorylated (lane 1, from left) and phosphorylated recombinant CovR protein (lane 2, from left) (**e**), CovR species were detected using an anti-CovR antibody and visualized using a fluorescently labeled secondary antibody (**e,g**). **f,h,** The percentages of CovR-P of total CovR protein were calculated using ImageJ. *n=3*. The data shown represent the means ± S.D. Statistical analysis was performed using an unpaired two-tailed t-test: *P <0.05 **P < 0.01 ***P<0.001 (**f,h**).

To further substantiate these findings, we quantified IL-8 cleavage using an ELISA assay for IL-8. The results clearly show that Asn stimulated ScpC activity production (Fig. 4c). As expected, the Δ*cov*S mutant produced a higher activity in the absence of Asn due to the basal increase in virulence gene expression. However, Asn-mediated ScpC increase in activity was still significant (Fig. 4c). However, in complete agreement with the data presented in (Extended Data Figs. 1c, d)., *cov*R*^-^* mutation fully activated ScpC activity, eliminating further increase by Asn (Fig. 4c). Furthermore, the Δ*asn*A *gln*P^-^ and *asn*B^-^ mutants lost or partially lost, respectively, their ability to produce ScpC in the presence of Asn, which was restored by their genetic complementation with WT genes (Fig. 4d). The somewhat higher ScpC activity of *asn*B^-^ in the absence and presence of Asn (Fig. 4d) probably is related to the lower level of its ATP content compared to the Δ*asn*A and *gln*P^-^ mutants (see Fig. 6).

Asn and LL-37 stimulated the production of ScpC in the GAS 5448 strain (Extended Data Fig. 4c) and in the GAS 854 strain (Extended Data Fig. 4d). Moreover, the Asn-mediated activation of ScpC in the GAS 854 strain was abolished in its *asn*A^-^ mutant derivative, suggesting that the Asn-mediated upregulation is ubiquitous among GAS M1T1 strains (Extended Data Figs. 4c, d).

Since the degree of phosphorylation of CovR regulates the expression of GAS virulence factors ^21, 28, 29^, we tested if Asn-mediated activation would also affect CovR∼P levels. To do so, we used Phos-Tag technology to quantify CovR phosphorylation^42^. First, we established that Asn does not reduce *in-vitro* phosphorylation of purified CovR by acetyl phosphate^43^ (Extended Data Fig. 4e). Then, we assessed CovR phosphorylation during the growth of S119 in CDM in the presence of Asn compared to unsupplemented CDM. As shown in Fig. 4e, the presence of Asn reduced the level of phosphorylated CovR, and some further reduction occurred in the presence of LL-37. Furthermore, Asn presence decreased the phosphorylation level of CovR in the S119Δ*cov*S mutant by more than 3-fold, demonstrating that CovS signaling is not required for Asn-mediated downregulation in CovR phosphorylation (Figs. 4e, f). To validate that the Asn-mediated decrease in phosphorylation of CovR was abolished in the Δ*asn*A mutant, we compared the phosphorylation of CovR of Δ*asn*A and the WT in the presence of Asn (Figs. 4e, f). It is apparent that the deletion of AsnA prevented the Asn-mediated decrease in CovR phosphorylation (Figs. 4g, h).

### Mutants deficient of *asn*A, *gln*P, and *asn*B are attenuated in a murine model of human NF

To test whether the mutants of AsnA, GlnPQ, and AsnB would have attenuated virulence during infection, we subjected Δ*asn*A, *gln*P^-^, and *asn*B^-^ to the sublethal soft tissue murine model of human NF that mimics the pathophysiology of the infection^44^. We enumerated colony-forming units (CFU) in soft tissue and spleen 2, 4, 6, and 8 days after infection (Figs. 5a-f). We also pictured the size of the developed lesions (Extended Data Figs. 5a-c) and measured the lesion area (Extended Data Figs. 5d-f). In addition, we determined the spleen weights (Extended Data Fig. 5g-i). Mice challenged with the indicated mutants were significantly attenuated compared to mice challenged with the WT strain S119 (Fig. 5a-f, and Extended Data Fig. 5a-i). The clearance rates of bacteria from the soft tissue and spleen and the decrease in lesion size and weight of the spleens of mice challenged with the mutants were more rapid and lighter than those of the WT S119 strain, respectively (Figs. 5a-f and Extended Data Figs. 5a-f). The *in vivo* data set concurs with the RT-qPCR determinations conducted for the indicated strains at OD = 0.7 in CDM in the absence and presence of Asn (Figs. 2 and 3) and the functional determinations of ScpA and ScpC activities (Fig. 4). Furthermore, an *asn*A^-^ mutant of GAS strain 854 was also attenuated in the same model (Extended Data Fig. 5j), demonstrating that Asn-mediated regulation of M1T1 virulence is ubiquitous among GAS M1T1 strains.

**Fig. 5.**
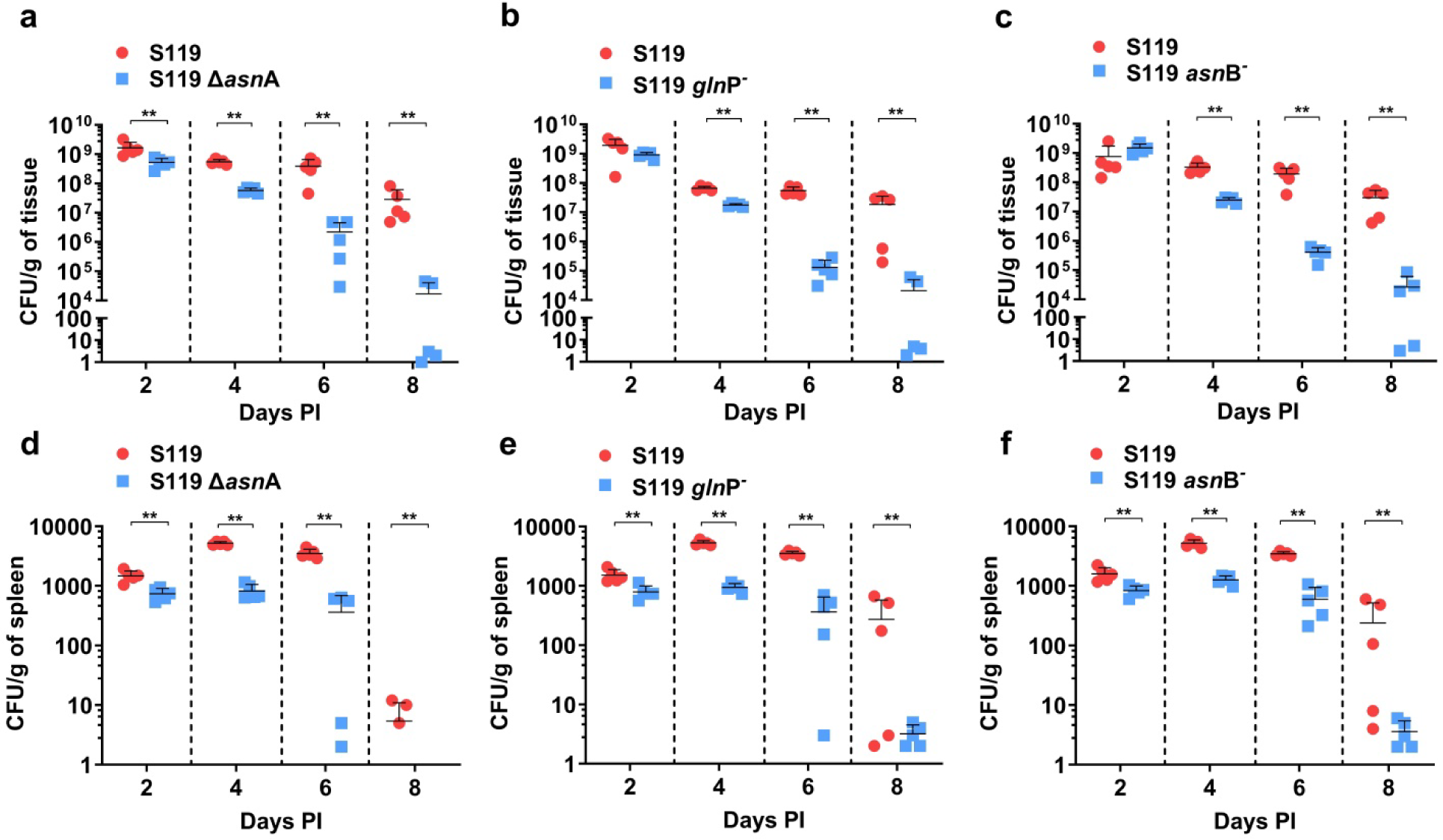
*asn*A, *gln*P, and *asn*B mutants are attenuated in the sublethal murine model of human NF. **a-c,** BALB/c mice were injected with a sub-lethal dose of GAS through a subcutaneous (SC) route. CFU counts per gram of soft tissue derived from mice (*n*=5) infected with S119 Δ*asn*A (**a**), S119 *gln*P^-^ (**b**), and S119 *asn*B^-^ (**c**) compared to the wild-type bacteria S119 were enumerated at indicated time points. **d-f,** CFU counts per gram of spleen derived from mice (*n=5*) after subcutaneous infection with S119 Δ*asn*A (**d**), S119 *gln*P^-^ (**e**), and S119 *asn*B^-^ (**f**) compared to S119, were determined at indicated time points. The data shown represent the means ± S.D. Statistical analysis was performed using the Mann-Whitney U test, *P< 0.05; **P < 0.01; ***P< 0.001 (**a-f**).

### Coupling of Metabolism to Virulence

GAS is a lactic acid bacterium that utilizes the glycolytic pathway but not the Krebs cycle due to the lack of critical enzymes^45^. To address the impact of Asn on the GAS metabolome, we assessed the level of intracellular and extracellular metabolites^36^. These experiments were conducted on the WT strain S119 and its derived *gln*P^-^ and *Asn*B^-^ mutants, grown in CDM to OD_600_ = 0.35 and 0.7 in the absence and presence of Asn (10 µg ml^-1^). The score plots of the probabilistic principal component analyses (PPCA) conducted on intracellular metabolites of all the samples at both optical densities for the indicated pairs of strains demonstrate that groups (clusters) were formed, thus statistically validating these measurements (Extended Data Figs 6a, b). The relative quantity of intracellular Asn in the S119 grown in the presence of Asn was significantly higher at OD_600_ = 0.35 than that of S119 grown in the absence of Asn or that of *gln*P^-^ mutant that cannot take up Asn (Fig. 6a). The relative amount of Asn of *Asn*B^-^ grown in CDM without Asn was almost as high as that of the S119 grown with Asn (Fig. 6a). Moreover, the relative Asn level of the *asn*B^-^ mutant grown in the presence of Asn was more than 80-fold higher than that of the WT grown under similar conditions, thus corroborating that the asparaginase activity of AsnB is very effective (Fig. 6a). At OD_600_ = 0.7, the relative Asn levels of S119 and *gln*P^-^ mutant grown in the absence and presence of Asn were comparable, whereas those of the *asn*B^-^ mutant were significantly higher under both conditions (Fig. 6a). Extracellular Asn level of showed that it was taken up entirely by the WT strain before the culture reached OD_600_ = 0.35 and remained negligible in the medium to OD_600_ = 0.7. Furthermore, no Asn was detected in the extracellular medium when GAS was grown in CDM without Asn, suggesting GAS does not release Asn to the medium during growth (Extended Data Fig. 6c). The *gln*P^-^ mutant did not transport Asn from the medium into the bacteria; thus, the relative Asn extracellular level remained constant and high (Extended Data Fig. 6c). The *asn*B^-^ mutant transported about 30% of the extracellular Asn at OD_600_ = 0.35; almost all of it was absorbed when the culture reached OD_600_ = 0.7 (Extended Data Fig. 6c). The reduced rate of Asn uptake of the *asn*B^-^ mutant compared to that of S119 probably results from a feedback inhibition exerted by the increased level of intracellular Asn (Extended Data Fig. 6c).

**Fig. 6.**
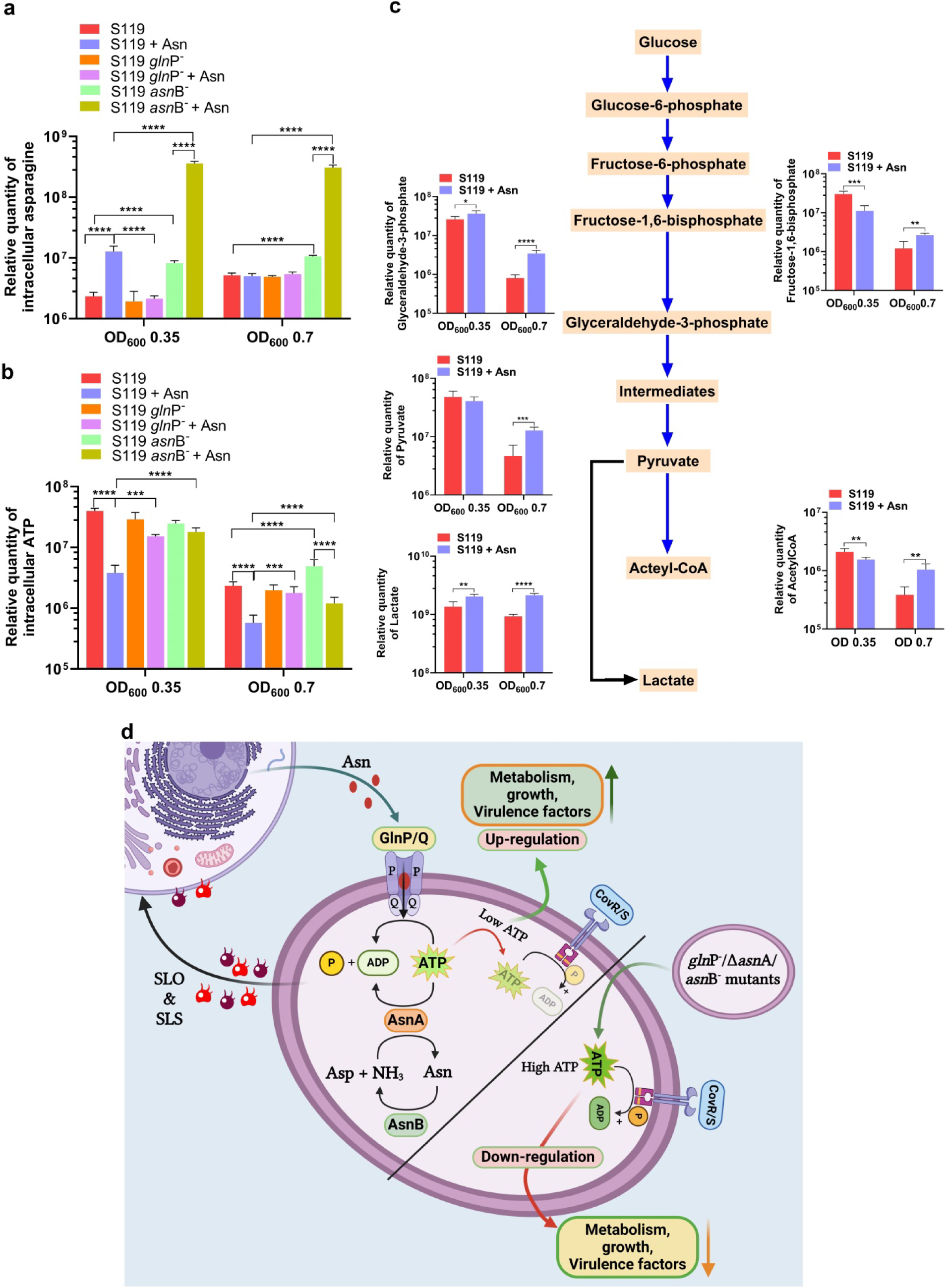
Coupling of Metabolism to Virulence. a,b, Intracellular metabolites. The relative amounts of intracellular Asn (**a**), and ATP (**b**), of the indicated strains grown in CDM or CDM-supplemented with Asn were determined. *n=5*. The data shown represent the means ± S.D. Statistical analysis was performed using an unpaired two-tailed t-test: *P <0.05 **P < 0.01 ***P<0.001 ****P< 0.0001. **c,** The measurements of some of the intracellular intermediates along the glycolytic metabolic pathway that converts glucose into pyruvate and lactate are presented and were determined for the S119 strain grown in CDM or CDM-supplemented with Asn to OD_600_ = 0.35 or 0.7. n=5. The data shown represent the means ± S.D. Statistical analysis was performed using an unpaired two-tailed t-test: *P< 0.05; **P< 0.01; ***P< 0.001, ****P< 0.0001 (**a-c**). **d,** Schematic representation of the mechanism controlling the regulation of CovR phosphorylation by modulating the intracellular ATP level in response to Asn in the WT S119 and its derived mutants.

To follow the metabolic status of S119 grown in the presence and absence of Asn at OD_600_ = 0.35 and 0. 7, we measured the relative levels of some of the intermediate metabolites along the metabolic pathways of GAS, leading to the formation of lactic acid from glucose^45^. We found that at OD_600_ = 0.35, the relationships between the levels of the denoted intermediates varied (Fig. 6b). However, at OD_600_ = 0.7, the relative levels of all metabolic intermediates were significantly higher for S119 grown in the presence of Asn, suggesting that Asn increased the rate of GAS metabolism (Fig. 6b). Indeed, the transcription of SP119_0416 encoding the glucose-6-phosphate isomerase was highly upregulated in the presence of Asn (Extended Data Figs. 1a, b). Furthermore, the transcriptions of the genes encoding the ATP synthase (SP119_0124, SP119_0125, SP119_0126, SP119_0127, SP119_0128 SP119_0129) were also significantly upregulated in the presence of Asn (GEO accession number GSE268517). Thus, it appears that Asn upregulates glucose metabolism to gain higher energy, which is committed to enhanced growth and increased gene expression.

The glucose levels in the extracellular CDM were similar for S119 and derived mutants grown in CDM in the presence and absence of Asn to OD_600_ = 0.35 and OD_600_ =0.7 (Extended Data Fig. 6d). Furthermore, the relative intracellular glucose levels at OD_600_ = 0.7 were comparable for the indicated strains, suggesting that glucose availability is not limiting for GAS metabolism and growth (Extended Data Fig. 6e).

Nevertheless, Asn strongly affects ATP levels and thus may constitute a limiting factor that connects virulence and growth with metabolism. In the presence of Asn, the level of ATP was the lowest for S119 compared to its *gln*P^-^ and *asn*B^-^-derived mutants both at OD_600_ = 0.35 and OD_600_ = 0.7 (Fig. 6c). As depicted in the schematic mechanism predicted in Fig. 6d, uptake of Asn from the host and intracellular production of Asn are ATP-requiring reactions strongly affecting the intracellular level of ATP, which, in turn, controls the degree of CovR phosphorylation. Thus, we suggest that Asn regulates GAS virulence by coupling metabolism to virulence (Fig. 6d).

## Discussion

GAS is a highly adapted and human-restricted pathogen. Therefore, it must derive its nutrition resources from the human host. GAS accomplishes this task by synthesizing various virulence factors capable of causing reversible minor damages to host cells at low doses, thus facilitating the attainment of essential nutrients. However, when produced in excess, these factors cause irreversible damage that can lead to lethality from invasive infections such as NF and STSS^8, 9^. Therefore, nutrient availability is expected to control virulence factor production tightly.

Here, we provide evidence that Asn regulates the expression of GAS virulence factors. This regulation is partly achieved by Asn feedback inhibition, which controls, together with CovR, the expression of SLO and SLS. Both toxins trigger ER stress and UPR, which upregulates the formation of Asn in infected host cells^17^. Thus, when Asn is in excess, the formation of extracellular Asn decreases, and *vice versa*. Another central circle of regulation is the link between the intracellular uptake of Asn by GlnPQ and the formation of Asn by AsnA and intracellular ATP levels. When GAS is cultured in CDM supplemented with 75 µM Asn to OD = 0.7, ATP but not glucose is limiting. AsnA and GlnPQ activities further consume the ATP. Consequently, virulence is increased because CovR is less phosphorylated. Most importantly, we demonstrate the physiological significance of this mechanism using the sublethal murine model of human soft tissue infection. This model allows us to follow the interaction between GAS and the host under balanced conditions between GAS virulence and the murine host defenses, leading to infection clearance of the WT strain within up to 15 days^44^.

What are the relationships between Asn-mediated gene regulation and GAS growth? Indeed, we could separate genetically between the Asn-mediated gene regulation and growth as the Δ*asn*A mutant grew well in the presence of Asn but could not regulate virulence genes and lacked the negative AsnA-feedback regulation. Nonetheless, Asn-mediated growth enhancement may involve regulatory circuits partially dependent on CovR phosphorylation or AsnA-feedback regulation. We previously reported that genes linked to replication, such as *pol*A, *lig*, *dan*X, and others, were upregulated by Asn^17^. The comparison of RNA-seq data of *cov*R, *cov*S, *asn*A, *gln*PQ, and *asn*B mutants should address the above question more directly.

Another open issue raised by our study is who is responsible for the CovS-independent phosphorylation of CovR, which is almost eliminated in CDM supplemented with Asn. It was suggested that acetyl phosphate plays a role in CovR phosphorylation *in vivo* for *Streptococcus mutans* that do not possess CovS natively^46^. The ATP levels in S119 grown in CDM in the presence of Asn are low and limiting (Fig. 6c). Acetyl phosphate concentration should be low under these conditions and would not reach the mM concentration required for phosphorylation. Thus, perhaps non-cognate sensor kinases or an orphan histidine-kinase regulator cause the CovR phosphorylation through a cross-talk mechanism, and they respond to changes in the intracellular ATP concentrations similarly.

As discussed above for GAS, Asn links metabolism to virulence under ATP-limiting conditions. Asn fulfills a similar role in cancer cells. The mitochondrial electron transport chain (ETC) activity is necessary for tumor growth, and its inhibition reveals anti-tumor treatment success when combined with targeted therapies^47^. For example, adding the Gram-negative asparaginase enzymes, such as Kidrolase, depletes Asn and, combined with metformin, is used as an anti-cancer treatment^6^. This treatment depletes intracellular Asn, increases ATF4 levels and thus the production of Asn-synthetase, and impairs mTOR complex I activity, essential for anabolic metabolism and cell growth^6^. We demonstrated that Kidrolase protects mice against peritoneal GAS infection and prevents bacterial proliferation in human blood, a hallmark of GAS infectivity^17^. In addition, as in the cancerous process, highly invasive GAS infections are usually accompanied by mutations in the CovR/S system that overproduces virulence factors, allowing GAS to become hypervirulent and overcome the host’s innate immunity defences^48^. In conclusion, both GAS infection and cancerous processes exploit the exquisite metabolic regulation of Asn under ATP-limiting conditions for their own benefit.

The mainstay of treatment for invasive GAS diseases is surgical debridement of infected tissues, prompt administration of intravenous antibiotics, and supportive care ^49,50^. Despite this, the associated mortality for invasive GAS diseases remains high, ranging from 23% to 35% in resource-rich settings^4, 9^. Furthermore, no safe and highly available vaccine against GAS exists^51^. Therefore, the necessity to develop effective novel treatments against GAS infections is self-evident. Based on this study, disconnecting the link between metabolism and virulence would attenuate GAS pathogenesis. Therefore, GlnPQ, AsnA, and AsnB-specific inhibitors should have therapeutic efficacy against invasive GAS diseases.

## Acknowledgments

We thank the core facility of the Faculty of Medicine at Hebrew University for conducting the RNA-seq analysis. We are indebted to Dr. Abed Nasereddin (Core Research Facility, Faculty of Medicine, The Hebrew University of Jerusalem) and Dr. Nevo Yuval (Unit of Bioinformatics, Faculty of Medicine) for their valuable contributions. We also greatly appreciate the support of Bella Agranovich and Ifat Abramovich from the Laura and Isaac Israel, Perlmutter Metabolomics Center, a part of the Biomedical Core Facility at the Technion, Israel Institute of Technology, Haifa, for their assistance in metabolomics experimental design, sample processing, and data analysis. This work was supported by the Israeli Ministry of Innovation, Science and Technology grant to EH (Grant number 0005663). The authors thank Prof. Herve Bercovier from the Department of Microbiology and Molecular Genetics, The Institute for Medical Research, Israel-Canada (IMRIC), Faculty of Medicine, The Hebrew University of Jerusalem, for reading this manuscript and for his valuable suggestions. Finally, we thank Prof. Samuel A Shelburne from the Department of Infectious Diseases, Infection Control, and Employee Health, The University of Texas MD Anderson Cancer Center, Houston, Texas, USA, for providing the necessary reagents and knowhow for conducting CovR phosphorylation assessments.

## Author contribution

E. H. conceived and supervised the study; A. S., A. A., and M. R. designed and conducted the experiments. A. S. and A.A. designed the figures and wrote the Materials and Method section. E. H. wrote the initial draft, which A. S., A. A., M. R., K. S. M, and N. H. edited. N. H. performed the experiment shown in Extended data Fig. 4e and provided all the reagents required for performing CovR-phosphorylation experiments, data described in Fig. 4 and in Extended data Fig. 4. K. S. M. supervised the study with E. H. and constructed some mutants. A. S. prepared the Figs. 3I and 6 and Extended data Fig. 6 and together with E. H., wrote the final version of this manuscript.

## Supplementary Data

**Extended Data Fig. 1.**
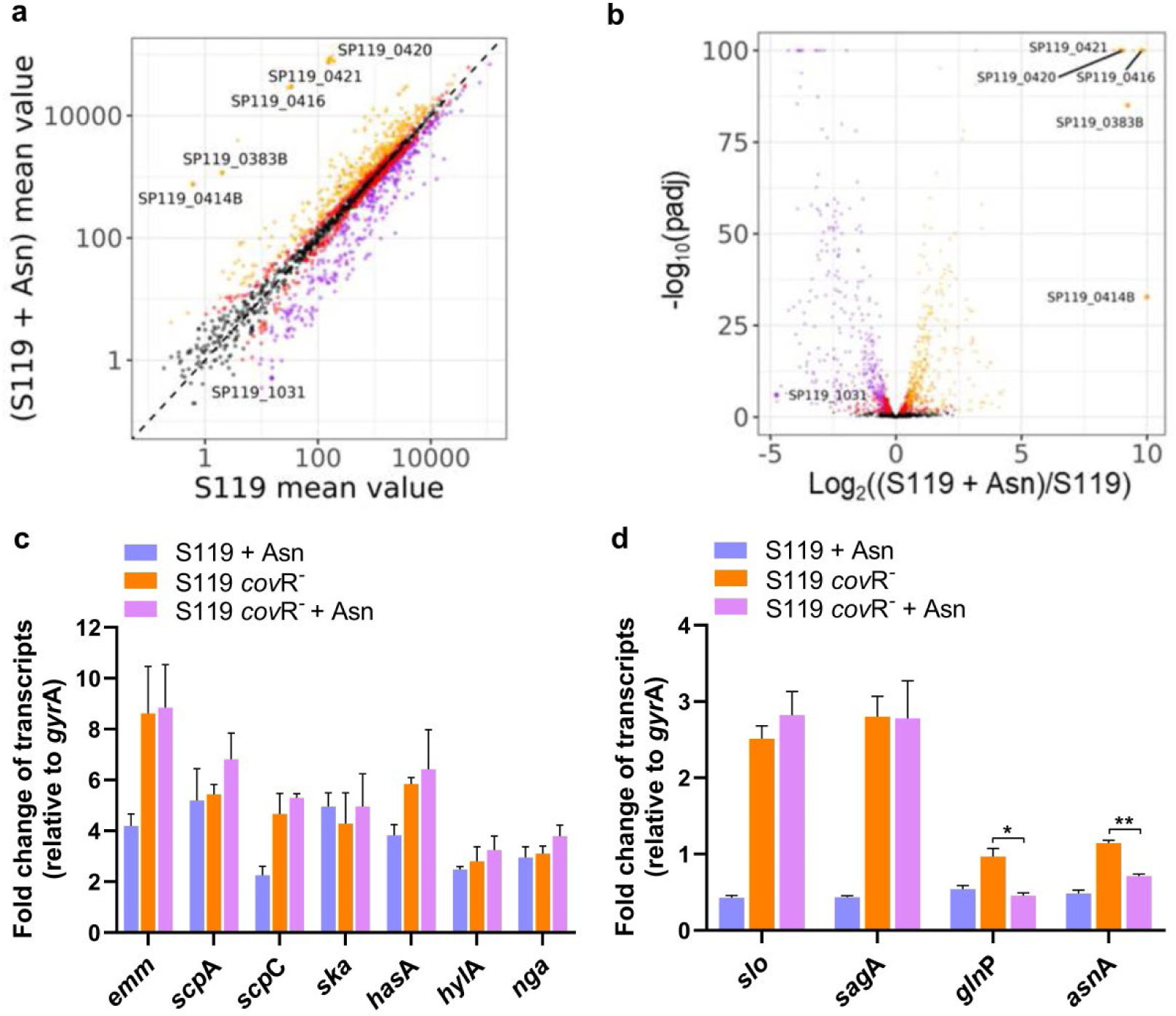
Asn regulates GAS transcriptome. (a,b) Visualization of mRNA-seq data. *n=4*. (a) A scatterplot matrix of the RNAseq data. Each dot represents a normalized mean value of the transcript number of the gene. (b) A Volcano plot of the same data set; each dot represents a gene with adjusted P < 0.05. (c,d) qRT-PCR determinations were performed for Set 1 (c), and Set 2 (d), genes on S119 and derived *cov*R^-^ mutant as in Fig. 1. Data shown represent the means ± S.D. Statistical analysis was performed using an unpaired two-tailed t-test; *P <0.05 **P < 0.01 (c,d).

**Extended Data Fig. 2.**
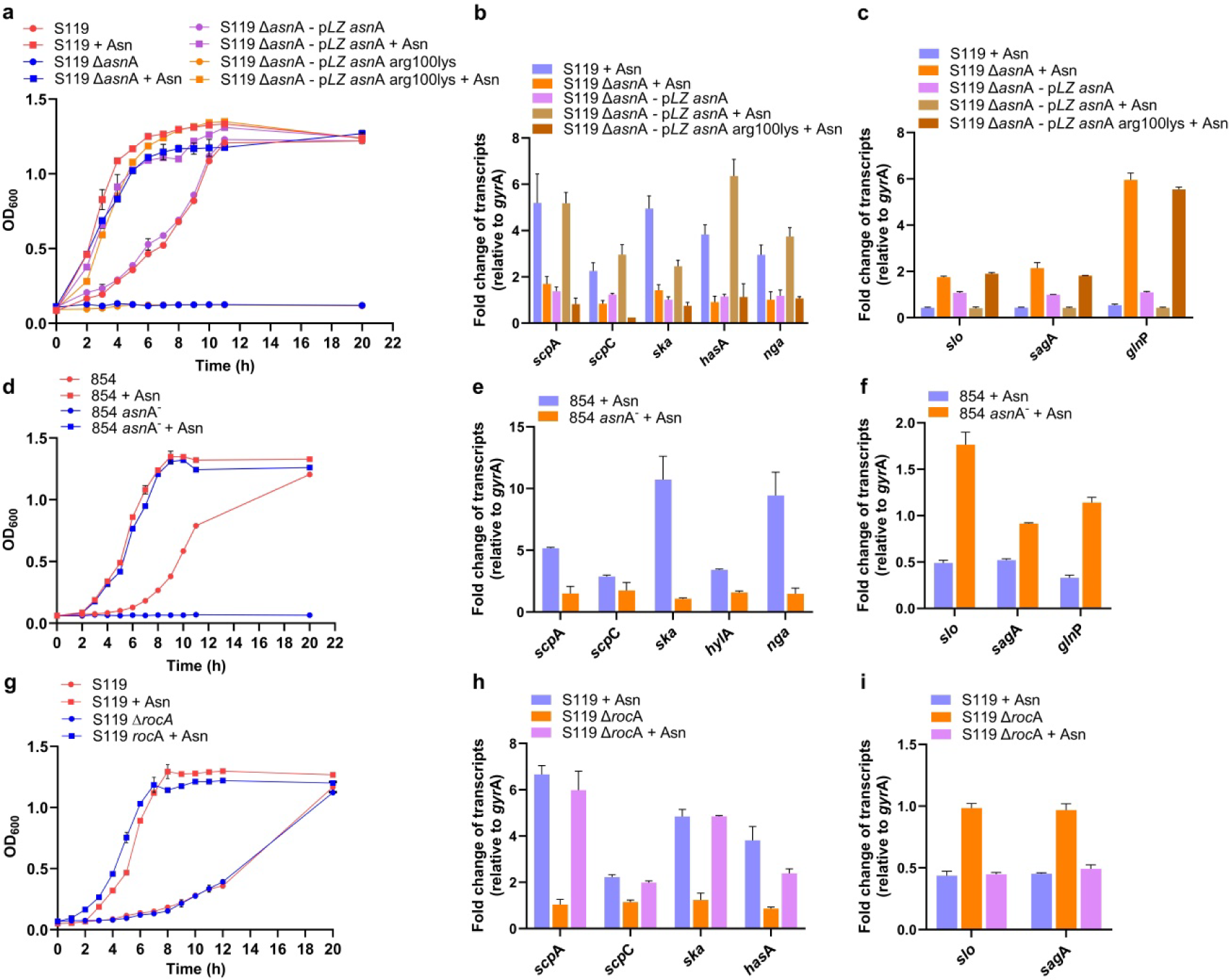
*asn*A is essential for Asn-mediated gene regulation. (a) The growth curves of stains S119, S119 Δ*asn*A, S119 Δ*asn*A*-*p*LZasn*A, and S119 Δ*asn*A-p*LZasn*A arg100lys. *n=2*. (b,c) qRT-PCR determinations for Set 1 (b) and Set 2 (c) were conducted as described in Fig. 2. *n=2*. (d) The growth of the GAS 854 strain and its *asn*A^-^derived mutant was monitored at indicated times. *n=2*. (e,f) qRT-PCR determinations for genes of Set 1 (e) and genes of Set 2 (f) were determined as described in Fig. 1. Statistical analysis was performed using an unpaired two-tailed t-test *P <0.05 **P < 0.01 ***P<0.001. **RocA is not involved in Asn-mediated effects.** The growth (g) and qRT-PCR determinations for Set 1 (h) slo and *sag*A, and (i) for S119 and its Δ*roc*A-derived mutant were performed as described above. The data shown represent the means ± S.D.

**Extended Data Fig. 3.**
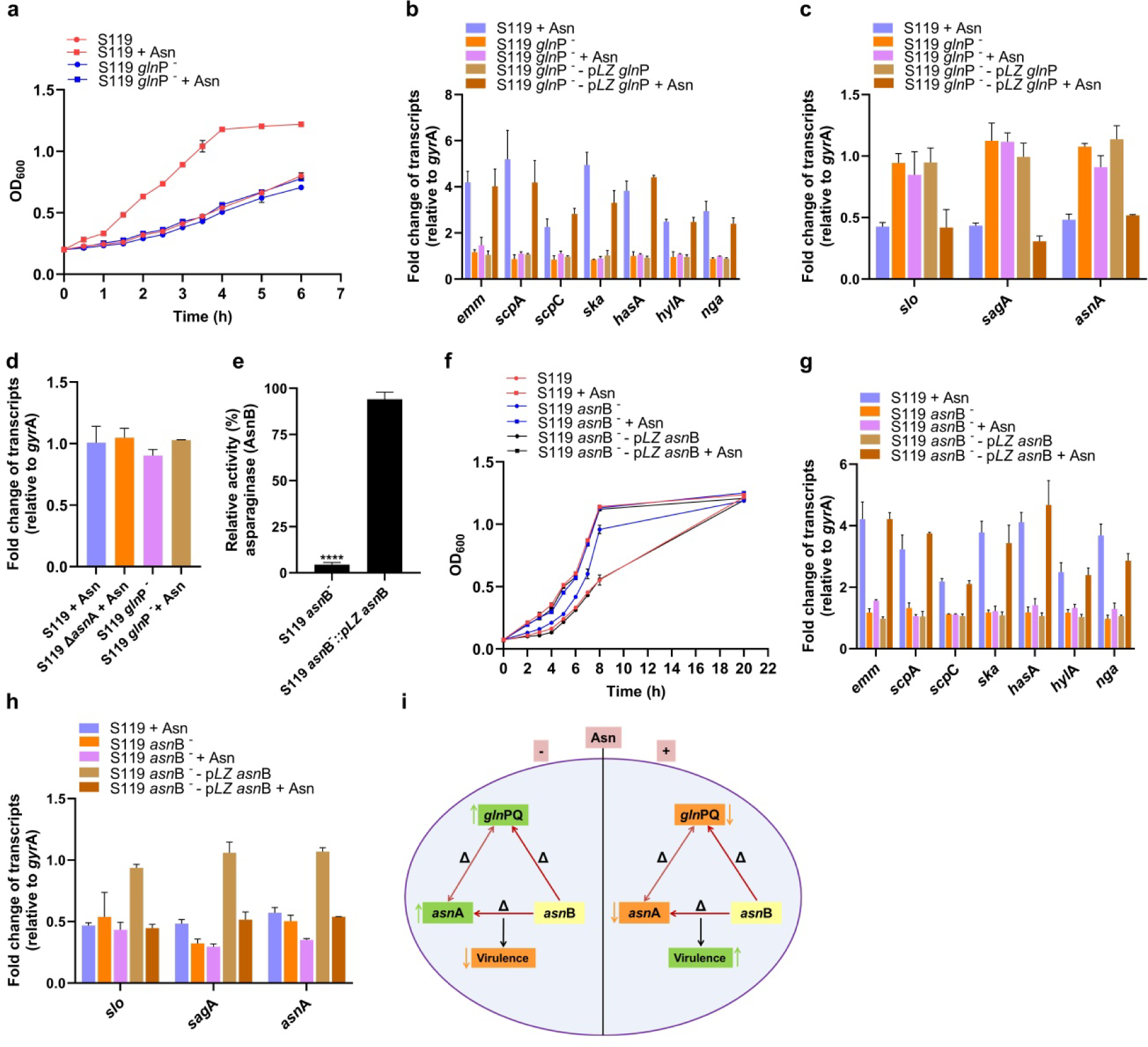
*gln*P and *asn*B are essential for Asn-mediated gene regulation. (a) The growth of S119 and S119 *gln*P^-^ in the absence or presence of Asn was determined (see Fig. 3b for the experimental details). *n=2*. (b,c) qRT-PCR determinations were conducted on Set 1 (b) or Set 2 of genes (c) using the indicated strains in CDM without or with Asn. *n=2*. (d) **The *asn*B transcript is unaffected by the presence or absence of Asn or by the deficiency of AsnA or GlnPQ activities.** qRT-PCR determinations of *asn*B transcript abundance were conducted as in (b,c). (e) Asparaginase activity (AsnB) was determined in the cell pellets of the indicated strains grown in THY. *n=2*. 100% represents the asparaginase activity of WT S119. (f) The growth of the indicated strains was monitored in CDM in the absence or presence of Asn (10 and 100 µg ml^-1^). *n=2*. (g,h) qRT-PCR determinations of Set 1 (g), and Set 2 (h), genes of the indicated strains grown in CDM without or with Asn. *n=2*. All qRT-PCR data (b,c,d,g,h) fold change was calculated by comparing with normalized transcript abundance in GAS S119 strain without Asn. The data shown represent the means ± S.D. (i) Graphical representation summarizes the orchestrated interplay between *asn*A, *gln*P, and *asn*B genes to maintain the balance of intracellular Asn in GAS.

**Extended Data Fig. 4.**
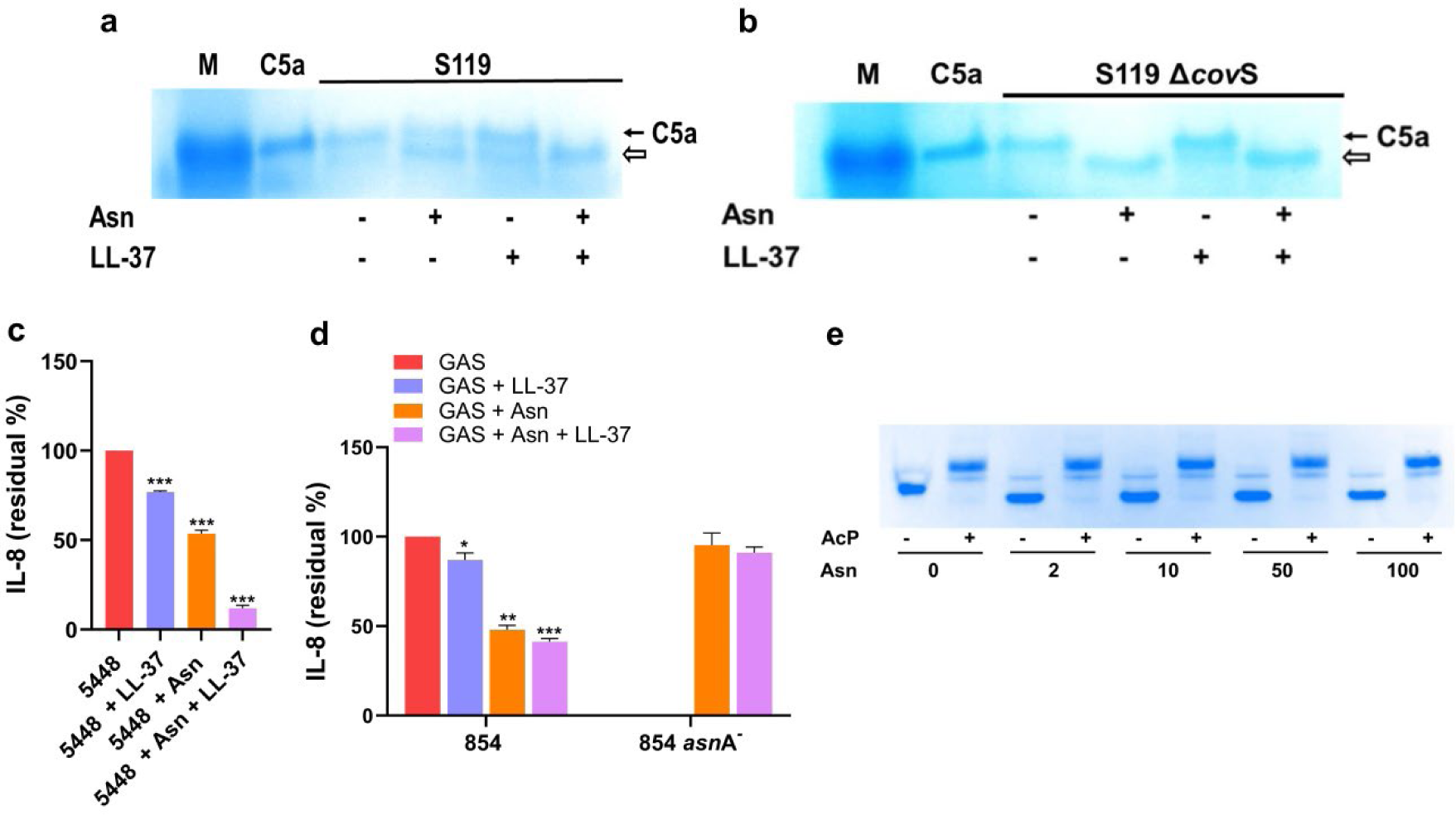
Functional assays and *in-vitro* phosphorylation of CovR. (a,b) Degradation of recombinant C5a by cultures supernatant containing the C5a-peptidase enzyme (ScpA). GAS was grown in CDM without or with Asn or/and LL-37. Supernatants of the indicated strains were incubated with recombinant human C5a, resolved on Tris-tricine gels, and visualized by Coomassie blue staining. M represents a marker, and an empty arrow is the cleaved C5a. The data are representative of 2 independent experiments. (c,d) Quantitation of IL-8 degradation by ScpC present in the supernatants of the indicated strains by ELISA. *n=2*. The data shown represent the means ± S.D. (e) **Asn does not affect *in-vitro* CovR phosphorylation**. Purified CovR was incubated in the absence (−) and presence (+) of acetyl phosphate (Ac-P) as a phosphate donor and the indicated concentrations of Asn (µg ml^-1^). The protein samples were resolved on Phos-tag SDS-PAGE gel and visualized.

**Extended Data Fig. 5.**
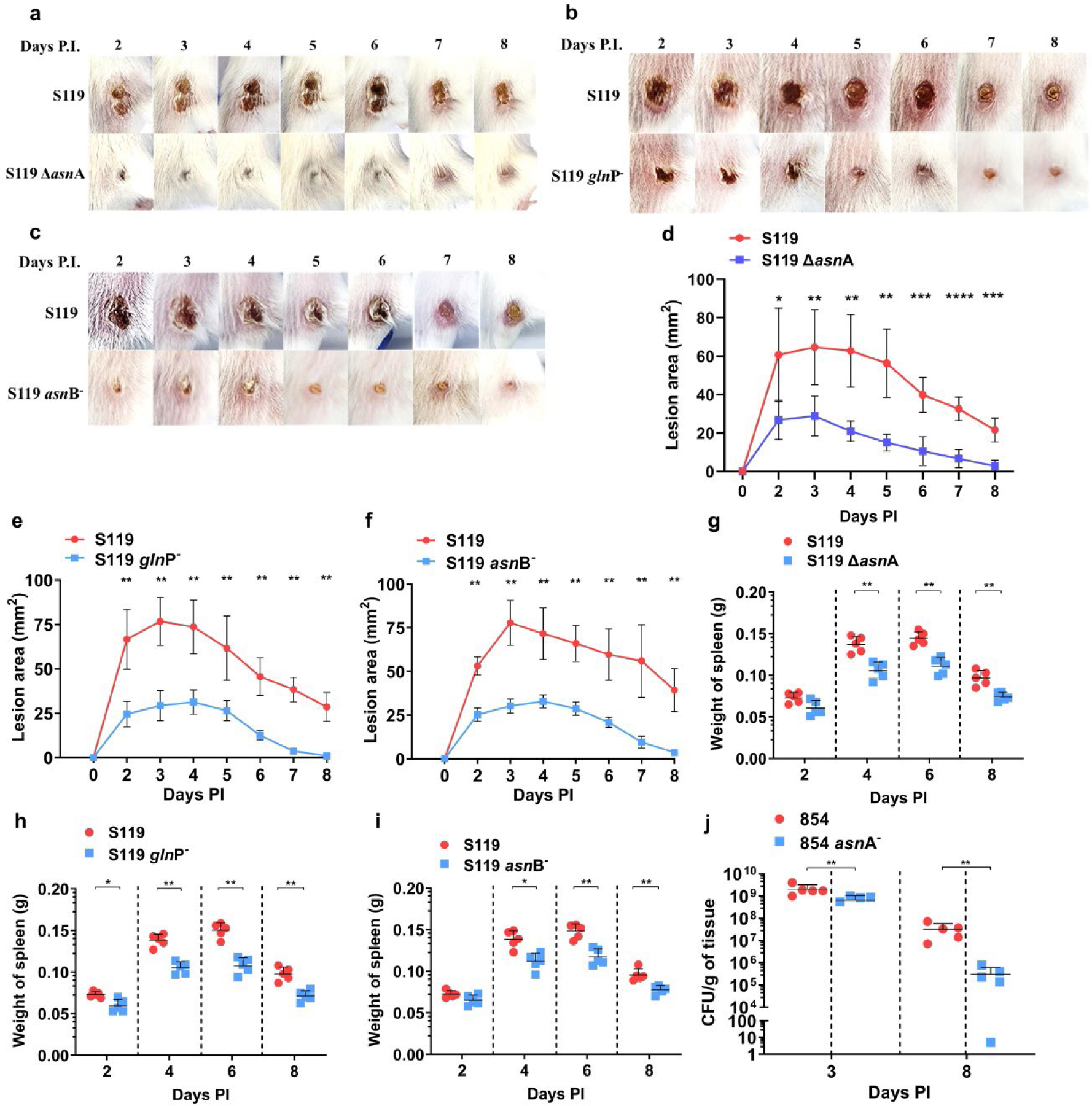
*asn*A, *gln*P, and *asn*B mutants are attenuated in the sublethal murine model of human NF. (a-c) Mice were injected subcutaneously, and representative images of lesion progression at indicated time points after S119 Δ*asn*A (a), S119 *gln*P^-^ (b), and S119 *asn*B^-^ (c), compared to WT S119 are shown. (d-f) Lesion areas of mice (*n*=5) infected S119 Δ*asn*A (d), S119 *gln*P^-^ (e), and S119 *asn*B^-^ (f) compared to S119 were determined at different time points post-infection. The data shown represent the means ± S.D. Statistical analysis was performed using the Mann-Whitney U test, *P < 0.05; **P < 0.01; ***P < 0.001 (d-f). **g-i** Spleenic weight of mice (*n*=5) infected subcutaneously with S119 Δ*asn*A (g), S119 *gln*P^-^ (h), and S119 *asn*B^-^ (i), in comparison to S119 was determined. The data shown represent the means ± S.D. The Mann–Whitney U test was used to determine the statistical significance, *P < 0.05; **P < 0.01; ***P < 0.001 (g-i). (j) **The deletion of *asn*A produces an attenuated mutant in GAS strain 854**. Mice were infected by subcutaneous, and CFU counts per gram of soft tissue (*n=5*) infected with 854 *asn*A^-^ compared to the wild type, 854 were enumerated at indicated time intervals. The data shown represent the means ± S.D. Statistical analysis was performed using the Mann-Whitney *U* test, *P< 0.05; **P< 0.01; ***P< 0.001

**Extended Data Fig. 6.**
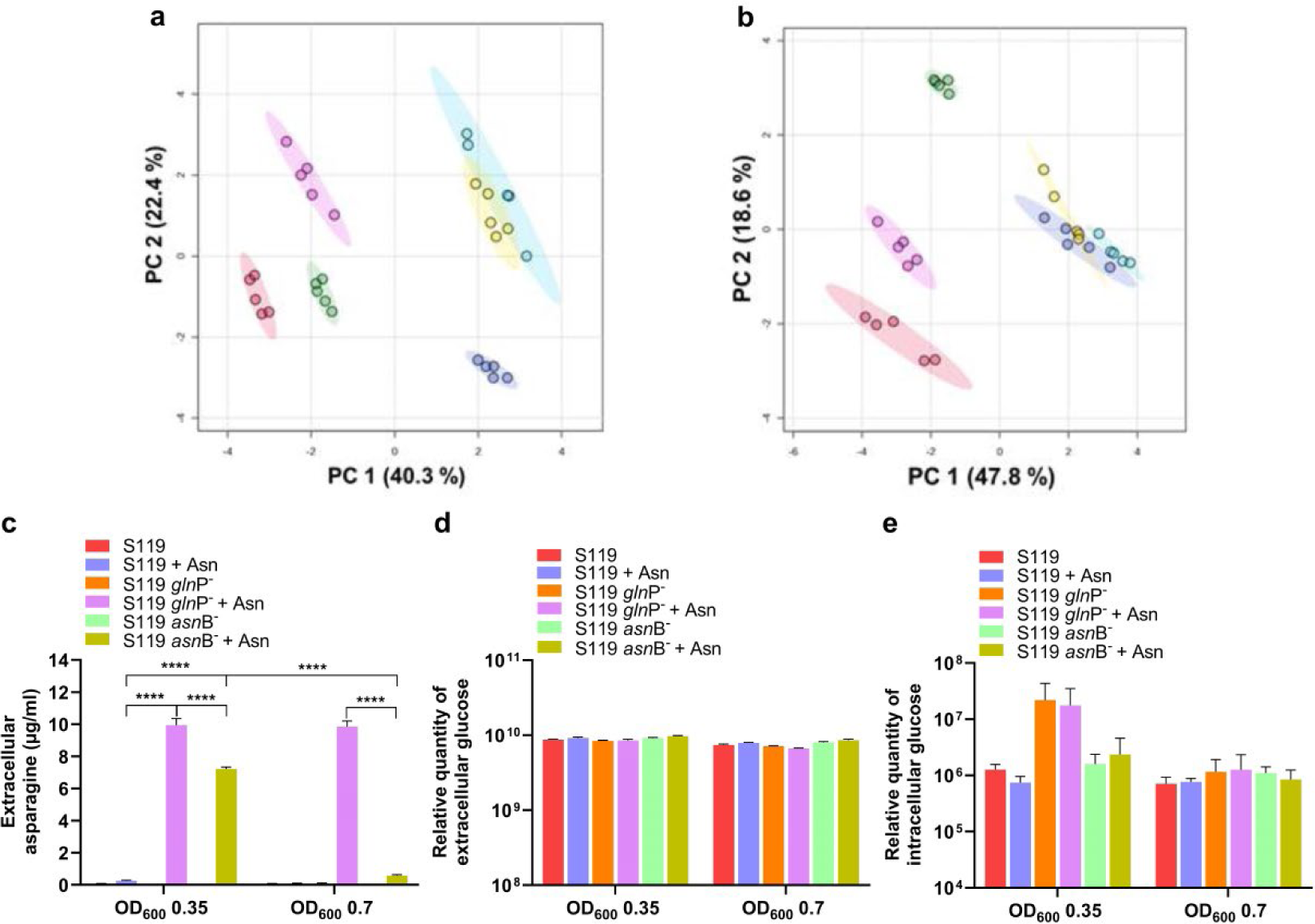
Coupling of Metabolism to Virulence. (a,b) The probabilistic principal component analyses (PCA) for the indicated pairs of strains grown in CDM or CDM supplemented with Asn to OD_600_ = 0.35 (a) or 0.7 (b). (c,d) Extracellular metabolite (c), the content of Asn was determined for the indicated strains grown in CDM or CDM-supplemented with Asn to OD_600_ = 0.35 or 0.7. *n=5*. The data shown represent the means ± S.D. Statistical analysis was performed using an unpaired two-tailed t-test. (d) The relative amount of glucose was determined for the indicated strains grown in CDM or CDM supplemented with Asn to OD_600_ = 0.35 or 0.7. *n=5*. (e) The relative amount of intracellular glucose was determined as above at OD_600_ = 0.35 or 0.7. *n=5*. The data shown represent the means ± S.D. Statistical analysis was performed using an unpaired two-tailed t-test. *P< 0.05; **P< 0.01; ***P< 0.001, ****P< 0.0001 (c,d).

## METHODS

### Bacterial culture

The GAS strains used in this study are represented in **Table S1**. GAS was cultured overnight without shaking in Todd-Hewitt broth supplemented with 0.2% yeast extract (THY) in sealed tubes at 37°C. When necessary, antibiotics were added at the final concentrations of 250 µg/ml for kanamycin (Km), 50 µg/ml for spectinomycin (Spec), or 1 µg/ml for erythromycin (Em). The following morning, overnight cultures were diluted 1:20 and grown in THY medium with appropriate antibiotic, when needed, to an early-log phase (OD600 of 0.3), washed, and resuspended in chemically defined medium (CDM) designed by van de Rijn and Kessler^1^. The growth rates of different bacterial strains were determined in CDM supplemented with or without asparagine (Asn) at different concentrations (2 to 100 µg/ml). 1 ml of freshly prepared CDM with bacteria (in the absence or presence of Asn and an appropriate antibiotic) was added to each well of a 24-well plate, and the plate was incubated at 37°C in a 5% CO_2_ atmosphere. The absorbance was measured at OD_600_ at regular time intervals. This method of culturing GAS in CDM was followed for all experiments.

### Construction of mutants

All mutants used in this study were generated using different primers (**Table S2**). **S119 Δ*cov*S** - a 718 bp fragment containing 107 bp of the end and 611 bp of the beginning of *cov*S was amplified with the primers *cov*RS-p3-F and *cov*R-R3 using S119 genomic DNA as a template. The PCR product was cloned into pGEM-T easy vector and then digested with *Eco*RV, and a Km-resistance gene (Ω*Km*) was cloned in this site. The resulting insert containing *cov*S upstream and downstream sequences separated by Ω*Km* gene was released from pGEM-T easy with *Eco*RI, treated with DNA Polymerase I, Large (Klenow) Fragment (NEB) for blunting the fragment, and then cloned into pJRS233 digested with *Eco*RI. The resulting plasmid, pJRS*cov*S-Ω*Km* was electroporated into strain S119 for knockout of *cov*S.

**S119 *cov*R**^-^ - a 296 bp fragment of *cov*R was amplified with the primers *cov*R-F2 and *cov*R-R3 using S119 gDNA as a template. The PCR product was cloned into pJRS233, which was digested with *Hin*dIII and *Bam*HI. The resulting plasmid, pJRS*cov*R, was electroporated into strain S119 for insertional inactivation of *cov*R. **S119 *asn*B^-^** - a 472 bp fragment of *asn*B was amplified with the primers *asn*B start- *Hin*dIII-L and *asn*Bstart-Pst using S119 gDNA as a template. The PCR product was cloned into pJRS233, and the resulting plasmid pJRS*asn*Bstart was electroporated into strain S119 for insertional inactivation of *asn*B.

**S119 *asn*B^-^ - p*L*Z *asn*B** - a 1071 bp fragment including the gene *asn*B and its upstream region was amplified with the primers *asn*B comp-F *Eco*RI and *asn*B comp-R-*Bam*HI using S119 gDNA as template. The PCR product was cloned into p*LZ*12, and the resulting plasmid p*LZasn*B comp was electroporated into strain S119 *asn*B^-^ strain to complement the *asn*B^-^ mutant.

**S119 *gln*P^-^** - a 671 bp fragment of *gln*P was amplified with the primers *gln*P-inact-F and *gln*P-inact-R using S119 gDNA as a template. The PCR product was cloned into pJRS233, and the resulting plasmid pJRS*gln*Pinact was electroporated into strain S119 for insertional inactivation of *gln*P.

**S119 *gln*P^-^** - **p*LZ gln*PQ -** A 3152 bp fragment *gln*PQ and its upstream region were amplified with the primers *gln*PQcomp-F-*Bam*HI and *gln*PQcomp-R-*Eco*RI using S119 gDNA as template. The PCR product was cloned into p*LZ*12, and the resulting plasmid p*LZgln*PQ comp was electroporated into strain S119 *gln*P^-^ to complement *gln*PQ.

**S119 Δ*asn*A** - a fragment containing 411 bp upstream of *asn*A, 380 bp downstream of *asn*A separated by Ω*Km* gene, was cloned into pJRS233, and the resulting plasmid pJRS*asn*A was electroporated into strain S119 for knockout of *asn*A. The 411 bp upstream fragment was amplified using the primers *asn*Aup-L-*Xba*I and *asn*Aup-R-*Sma*I and GAS JS95 gDNA as a template. The 380 bp downstream fragment was amplified using the primers *asn*ADown-R-*Ps*tI and *asn*ADown-L-*Sma*I using JS95 gDNA as a template.

**S119 Δ*asn*A-p*LZ asn*A** - a 1703 bp fragment including the gene *asn*A and its upstream and downstream regions was amplified using the primers *asn*Aup-L-*Xba*I and *asn*ADown–R using S119 genomic DNA as a template. The PCR product was cloned into p*LZ*12, and the resulting plasmid p*LZasn*A comp was electroporated into strain S119 Δ*asn*A for complementation of *asn*A.

**S119 Δ*asn*A-p*L*Z *asn*A R100K** - p*L*Z *asn*A R100K was constructed by amplifying p*L*Z*asn*A with 2 sets of primers: Gibson-p*LZ* km-F + Gibson-p*LZ asn*A-Arg100-Lys-R and Gibson-p*LZ* Km-R + Gibson-p*LZ asn*A-Arg100-Lys-F, using p*L*Z*asn*A as a template. The amplified PCR products were ligated using a Gibson assembly kit (NEB, USA). The resulting plasmid p*L*Z *asn*A R100K was electroporated into strain S119 Δ*asn*A for expression of mutated *asn*A.

**S119 Δ*roc*A** - a 3298 bp fragment containing *roc*A flanked by its upstream and downstream sequences was PCR amplified with the primers *roc*A up-F and *roc*A down-R using S119 gDNA as a template and was cloned into pGEM®-T Easy Vector. The resulting plasmid pGEM *roc*A was digested with *Xba*I and *Hpa*I, releasing a 971 bp fragment of *roc*A, and a km-resistance gene was inserted. The resulting fragment containing *roc*A upstream and downstream sequences separated by Ω*Km* gene was cloned into pJRS233, and the resulting plasmid pJRSR*roc*A was electroporated into strain S119 for knocking out *roc*A.

### Extraction of RNA from GAS and qRT-PCR determinations

GAS was grown in CDM in the absence or presence of Asn (10 or 100 µg/ml) or 300 nM LL-37 in 24-well plates at 37°C in 5% CO_2_ atmosphere, and samples were collected at late log phase (OD_600_ = 0.7). Samples were centrifuged at 5,000 x g for 10 min at 4°C. Supernatants were discarded, and the pellet was resuspended in 1 ml of RNA protect bacteria reagent (Qiagen). Then, samples were centrifuged for 1 min, the supernatant was discarded, and the pellet was stored at -80°C until use. Next, the pellet was resuspended in 350 µl of RNA1 solution (100 mM Tris pH 7.0, 1 mM EDTA, and 25% glucose), 70 µl of fresh lysozyme (9 mg/ml), and 7 µl of mutanolysin (10,000 U/µl) were added, and the samples were incubated for 5 min at 37°C. After incubation, samples were centrifuged at 14,400 x g for 2 min at 4°C, and supernatants were discarded. The pellets were resuspended in 300 µl of RNA2 solution (20 mM sodium acetate pH 5.5, 1 mM EDTA, and 0.5% SDS). A total of 300 µl of water-saturated bio-phenol was added. The mixtures were incubated for 6 min at 65°C while vortexing every minute for 10 sec. Then, samples were centrifuged at room temperature for 5 min at 14,400 x g. The upper phase was removed to a new tube, an equal volume of phenol and 1 volume of ethanol (95-100%) were added, and the suspensions were transferred to the Zymo-spin IIC column. According to the manufacturer’s instructions, RNA purification was conducted using the Direct-zol RNA miniprep kit (Zymo Research). The RNA concentration and purity were evaluated using NanoDrop One (Thermo Scientific), and intactness was determined following visualization on 1% agarose gel. RNA was treated with RQ1 DNase (Promega) according to the manufacturer’s instructions to avoid genomic DNA contamination, and total DNA degradation was verified by PCR using the primers *sra*-f and *sra*-r. M-MLV Reverse Transcriptase (Promega) was used for reverse transcription according to the manufacturer’s instructions, and cDNA synthesis was verified by PCR using the same set of primers. For real-time PCR, cDNA was diluted, and quantitative PCR was performed on Mic qPCR Cycler (Bio-Molecular Systems) using the 2X Tamix Fast SyGreen Mix Hi-ROX (Tamar Laboratory Supplies Ltd). The primers used are listed in **Table S2**. Each target gene’s expression amounts were normalized to *gyr*A and analyzed using the 2-ΔΔC_T_ method (Livak et al., 2001). Experiments were performed in at least duplicates from two independent cDNA preparations. The standard deviation was determined using Prism 10 (GraphPad Software).

### RNA-Seq and data analysis

For RNA-seq, GAS was cultured in CDM in the absence or presence of Asn (10 µg/ml) in 24-well plates at 37°C in a 5% CO_2_ atmosphere, and samples were collected once the OD_600_ reached 0.7. Samples were centrifuged at 5,000 x g for 10 min at 4°C. RNA was extracted and stored at -80^0^C until further use. The RNA samples were treated with the RQ1 DNase (Promega) according to the manufacturer’s instructions to avoid genomic DNA contamination. Ribosomal RNA removal was achieved using the Pan-Bacteria (RNA-Seq) riboPool^TM^ kit (siTOOLs BioTech, Germany). Sample quality was assessed using a 2100 Bioanalyzer (Agilent), and sample quantity was determined using a NanoDrop 8000 spectrophotometer (Thermo Scientific). According to the manufacturer’s recommendations, the RNA-seq directional libraries were generated for 1 μg rRNA-depleted RNA using the KAPA Stranded mRNA-Seq Kit (Kapa Biosystems, KK8421). The rRNA-depleted RNA was fragmented, followed by cDNA synthesis using random priming. The cDNA strands were purified using the AMPure System (Agencourt), and A-tailing modification was performed. Then, adapter ligation was done using Uniquely Dual Indexed (UDI) primers, the library was amplified, and molarity was determined using the Qubit system (Qubit® DNA HS Assay kit, Invitrogen), and the TapeStation system, using DNA HS ScreenTape kit (Agilent Technologies). A 76 bp single-read DNA sequencing was performed at the Core Research Facility, Faculty of Medicine, The Hebrew University of Jerusalem, Israel, using the Illumina Nextseq500 platform. Data were generated in the standard Sanger FastQ format, and raw reads have been deposited under BioProject GSE268517 with the Sequence Read Archive (SRA) at the National Center for Biotechnology Institute.

The NextSeq basecalls files were converted to fastq files using the bcl2fastq program with default parameters (without trimming or filtering applied at this stage). Raw reads (fastq files) were inspected for quality issues with FastQC. Following that, the reads were quality-trimmed with Cutadapt, using a quality threshold of 32 for both ends, poly-G sequences (NextSeq’s no signal) and adapter sequences were removed from the 3’ end, and poly-T stretches were removed from the 5’ end (being the reverse-complement of poly-A tails). The Cutadapt parameters included using a minimal overlap of 1, allowing for read wildcards, and filtering out reads that became shorter than 15 nt. Finally, low-quality reads were filtered out using fastq_quality_filter, with a quality threshold of 20 at 90 percent or more of the read’s positions. The processed reads were aligned to the reference genome of *Streptococcus pyogenes* strain S119 (GCA_900608505.1, Genebank) with bowtie2 version 2.3.4.3. Quantification was done with htseq-count. Strand information was set to ’reverse.’ All counts of none CDS (ncRNA, antisense_RNA, rRNA, tRNA, SRP_RNA, RNase_P_RNA) genes were removed from the count’s Table and subjected to differential expression analysis. All the processed reads were aligned to the *S. pyogenes* strain S119 genomic rRNA sequence using bowtie2 version 2.3.4.3 to measure the amount of rRNA-originating reads. Normalization and differential expression analysis were done with the DESeq2 package. Genes with a sum of counts less than 10 over all samples were filtered out, and then size factors and dispersion were calculated. Normalized counts were used for several quality control assays, such as counts distributions and principal component analysis, which were calculated and visualized in R. Pair-wise comparisons were tested with default parameters, except not using the independent filtering algorithm. The significance threshold was taken as p_adj_<0.1 (default). In order to reduce the number of false positive results, significant genes were further filtered. The criteria for filtering included a requirement for a significant log_2_ foldchange, i.e., baseMean above 5 and an absolute change more significant than 5/sqrt(baseMean)^0.7 + 0.5 (for highly expressed genes. This requirement means a fold-change of at least 1.5, while genes with a significantly reduced expression would need a 7-fold change to pass this filtering. While the genes that passed the default (p_adj_<0.1) filter are termed "Sig," the sub-set of genes that also passed the additional filter is termed "Best." Finally, results were combined with gene details from Genbank’s feature Table.

### ScpC cleavage assay

GAS was cultured in CDM in the absence or presence of Asn (10 µg/ml) or 300 nM LL-37 in 24-well plates at 37°C in 5% CO_2_ atmosphere, and samples were collected at late log phase (OD_600_ of 0.7). Samples were centrifuged at 5,000 x g for 10 min at 4°C, and supernatants were collected. The cell-free supernatants were incubated with an equal volume of 1 mg/ml of recombinant human IL-8 (R&D Systems, USA) at 37°C for 2h. The samples were heated at 100°C for 5 min with 4x Tricine loading buffer (0.4 M Tris HCl pH 6.8, 80% glycerol, 4% SDS, and 0.08% Coomassie blue) to stop the reaction. A total volume of 25 µl samples was loaded on precast 16.5% Mini-PROTEAN® Tris-Tricine gels using the Mini-PROTEAN gel apparatus (Bio-Rad). A total of 0.5 µg of purified recombinant IL-8 (R&D Systems, USA) and protein marker (Low-range Rainbow molecular weight marker, DUTSCHER) was loaded as a control. The samples were run on a low constant voltage at 4°C for 6 to 8 h in Tris Tricine SDS Running Buffer (Bio-Rad, USA). The peptides were detected by Instant Blue (Expedeon Inc.). The gels were de-stained with distilled water until the bands were visible.

### Cleavage of ScpA

GAS was grown in CDM in the absence or presence of Asn (10 µg/ml) or 300 nM LL-37 in 24-well plates at 37°C in 5% CO_2_ atmosphere, and samples were collected at late log phase (OD_600_ of 0.7). Samples were centrifuged at 5,000 x g for 10 min at 4°C, and supernatants were collected. The cell-free supernatants were incubated with 1 mg/ml of purified recombinant human complement component C5a (R&D Systems, USA) at 37°C for 2 h. The samples were heated at 100°C for 5 min with 4x Tricine loading buffer (0.4 M Tris HCl pH 6.8, 80% glycerol, 4% SDS, and 0.08% Coomassie blue) to stop the reaction. A total volume of 25 µl samples was loaded on precast 16.5% Tris-Tricine gels using the Mini-PROTEANgel apparatus (Bio-Rad). A total of 0.5 µg of purified recombinant C5a (R&D Systems, USA) and protein marker (Low-range Rainbow molecular weight marker, DUTSCHER) was loaded as a control. The samples were run on a low constant voltage at 4°C for 6 to 8 h in Tris Tricine SDS Running Buffer (Bio-Rad, USA). The peptides were detected by Instant Blue (Expedeon Inc.). The gels were de-stained with distilled water until the bands were visible.

### ELISA-based assessment of ScpC expression by performing IL-8 degradation assays

GAS was grown in CDM in the absence or presence of Asn (10 or 100 µg/ml) or 300 nM LL-37 in 24-well plates at 37°C in 5% CO_2_ atmosphere, and samples were collected at late log phase (OD_600_ of 0.7-0.8). Samples were centrifuged at 5,000 x g for 10 min at 4°C, and supernatants were collected. The cell-free supernatants were incubated with 1 ng/ml of recombinant human IL-8 (R&D Systems, USA) at 37°C for 2 h. The samples were heated at 100°C for 2 min; the reaction was stopped. IL-8 cleavage was represented as the residual amount of IL-8 in the samples, estimated using the Human IL-8 Quantikine ELISA kit (R&D Systems, USA).

### Extraction of protein from GAS pellets

Bacterial cultures were grown overnight in THY media and harvested the following day. After washing in sterile PBS, it was suspended in CDM without/with Asn (10 µg/ml). The bacterial suspension was distributed in 24-well plates and incubated in 5% CO_2_ at 37°C until the OD_600_ reached 0.6 to 0.7. All further steps were performed at 4°C. Bacterial cultures were collected, centrifuged, and washed with ice-cold sterile PBS. For extraction, the pellet was suspended in ice-cold lysis buffer [20 mM Tris-HCL pH 8 with 10 U mutanolysin (Sigma Aldrich), cOmplete^™^, EDTA-free Protease Inhibitor Cocktail (Roche Molecular Diagnostic, USA), and PhosSTOP™ (Roche Molecular Diagnostic, USA) in PBS]. The bacterial suspension was added to lysing matrix tube B (0.1mm) (MP Biomedicals) and homogenized by using 6500 rpm for 30 sec (MagNA Lyser, Roche Diagnostics). This step was repeated thrice with a time interval of 1 min. The lysates were centrifuged at 12000g for 5 min at 4°C, and the supernatants were collected and stored at -80°C until further use. The total protein was quantified by Bradford’s assay and equalized in each sample with sterile DDW.

### Phos-tag western blotting to measure phosphorylation of CovR

Before electrophoresis, a total of 67 µl of 4x Laemmli Sample Buffer (900 µl of Laemmli Sample Buffer + 100 µl of ß-mercaptoethanol) was added to 200 µl of protein sample. For resolving phosphorylated and non-phosphorylated proteins, a total of 10 µg of protein sample was added to each well of Phos-tag SuperSep Phos-tag Gels (Zn^2+^ and 12.5% with 50 µM of phostags, 17 wells of 30 µl capacity from Wako). Recombinant CovR was purified as described^2^ and served as control. The *in vitro* phosphorylation of recombinant CovR protein was performed according to previously published work^3^. In brief, purified CovR protein was incubated for 2 h in 50 mM Tris-HCl, pH 8.0, 10 mM MgCl_2_, 3 mM dithiothreitol, and 32 mM acetyl phosphate. The gel was run in Tris-Glycine Running Buffer (Bio-Rad, USA) at low voltage (50 V) and constant current in a 4°C until the protein marker (BLUeye Prestained Protein Ladder, Sigma-Aldrich) front reached the end of the gel. The gel was treated with blotting buffer [10x SDS PAGE buffer 50mL (5 of 1X) + MetOH 200 ml (2) + adjusted to total volume of 1 liter (milliQ)] and supplemented with 10 mM EDTA. The gel was shaken for 10 min 3 times to remove all Zn^2+^ from the gel.

Then, the gel was rewashed with the blotting buffer without EDTA. The blotting was performed using the Trans-Blot Turbo Transfer Pack (Bio-Rad) at a constant voltage of 10 V for 1 h. The membrane was blocked overnight at 4°C in a solution containing PBST supplemented with 5% skim milk. The membrane was washed in PBST and placed in a suspension of rabbit anti-CovR antibody in PBST (1:5000) for 1 h at room temperature. The membrane was washed 4 times in fresh PBST and treated with fluorescently labeled Goat Anti-Rabbit IgG StarBright Blue 700 antibody (1:5000) in 10 ml of PBST buffer containing 1% skim milk at room temperature for 2 h. Finally, the membrane was washed 4x with PBST. The fluorescently labeled proteins were visualized at the wavelength of 470 nm by using the Biorad GelDoc system. The relative percentage of phosphorylated CovR was calculated using ImageJ software (open source, developed by NIH, USA).

### Asparaginase assay

We used the Nesslerization method to determine the intracellular L-asparaginase activity in whole-cell suspension [Roy et al., 2019]. Overnight-grown GAS was inoculated into fresh THY media and incubated at 37°C until OD_600_ reached 0.7. Cells were harvested by centrifugation at 14000 rpm for 5 min, and the cell pellet was washed twice with sterile saline. The washed pellet obtained from 50 ml culture was suspended in 190 µl of sterile PBS, and 10 µl of pure PlyC phage protein (1 µg/µl) was added to make up the total volume of 200 µl. The lysis was completed at 37°C in 5 minutes. An equal volume (100 µl of each) of lysate and reaction mixture (50 mM L-Asn and 100 mM Tris-HCl, pH 8.0) was mixed and incubated at 37°C for 10 min. Afterwards, the reaction was terminated by adding 20 µL trichloroacetic acid (1.5 M). After centrifugation at 14000 rpm for 5 min, 15 µL Nessler’s reagent was added to the supernatant, and the absorbance was measured at 436 nm.

### Metabolomics

For time-dependent Asn uptake kinetics, bacterial cultures (S119 and S119 *gln*P^-^) were grown overnight in THY media and harvested the following day. After washing in sterile PBS, it was suspended in CDM without Asn until the OD_600_ reached 0.2. Asn was added (final concentration of 10 µg/ml) in culture, and the bacterial suspension was distributed in 24-well plates and incubated in 5% CO_2_ at 37°C for 1h. The samples were collected at different time points (5, 30, and 60 min), and a total of 20 µl of sterile filtered supernatant was mixed with 980 µl of chilled extraction solution (50% methanol, 30% acetonitrile, 20% water) by brief vortexing and stored at -80°C until further analysis.

For extracellular and intracellular metabolites detection, overnight grown bacterial culture (S119, S119 *gln*P^-^, S119 *asn*B^-^) as mentioned above were washed and inoculated in fresh CDM without or with Asn (10 µg/ml). The bacterial suspension was distributed in 24-well plates and incubated in 5% CO_2_ at 37°C until OD_600_ reached 0.35 or 0.7. For extracellular metabolite detection, 0.35 and 0.7 OD_600_ samples were collected at OD_600_ of 0.35 and 0.7, and a total of 20 µl of sterile filtered supernatant was mixed with a total of 980 µl of chilled extraction solution as above until further analysis. For intracellular metabolite detection, a total 2 ml of culture was collected centrifuged, and pellets were washed with sterile PBS and suspended in 95 µl of sterile PBS. A total of 5 µl of pure PlyC phage protein (1µg/µl) was added to make up the total volume of 100 µl. The lysis was completed at 37°C in 5 minutes. A total of 30 µl of lysate was mixed with a total of 300 µl of chilled extraction solution as above. All samples were stored at -80°C until further analysis. The Bradford assay was used to estimate protein in intracellular samples. The protein quantity in each sample was used to normalize the quantity of detected metabolites. All experiments were conducted in five biological replicates (n=5).

LC-MS metabolomics analysis was performed as described previously (PMID: 26358905) with slight changes for polar metabolite detection. ThermoFisher Scientific Vanquish ultra-high-performance liquid chromatography (UHPLC) system coupled to Exploris 240 Orbitrap Mass Spectrometer (ThermoFisher Scientific) was used with a resolution of 120,000 at 200 mass/charge ratio (m/z), electrospray ionization, and polarity switching mode to enable both positive and negative ions across a mass range of 67 to 1000 m/z. HPLC setup consisted of a ZIC pHILIC column (SeQuant; 150 mm x 2.1 mm, 5µm). 5 µL of biological extracts were injected, and the compounds were separated using a mobile phase gradient of 15 min, starting at 20% aqueous (20 mM ammonium carbonate (ThermoFisher Scientific, 10785511) adjusted to pH = 9.2, with 0.1% of 25% ammonium hydroxide (ThermoFisher Scientific, 15547049) and 80% acetonitrile. The gradient was terminated with 20% acetonitrile. Flow rate and column temperature were maintained at 0.2 mL/min and 45°C, respectively, for a total run time of 27 minutes. All metabolites were detected using a mass accuracy below 5 ppm. Xcalibur (ThermoFisher Scientific) was used to acquire data.

Skyline Daily (PMID: 31984744) version 23.1.1.503 was used for analysis. Instrument settings were set to orbitrap, and the resolution was set at 120,000 at 200 m/z. The transition list containing the compound names, accurate masses, explicit retention times with a 2-minute retention time window, and the polarity was used to generate extracted ion chromatograms for each compound, and chromatographic peaks were inspected and integrated. Relative quantification between sample groups was performed using the area of the signal. Metabolite AutoPlotter 2.6 (PMID: 32670572) and Metaboanalyst (PMID: 34019663) were used for data visualization.

### Animals

3-to 4-week-old female BALB/c OlaHsd mice weighing 10-12 g were obtained from ENVIGO RMS (Israel Ltd.). Following the Hebrew University of Jerusalem’s ethical guidelines, all procedures were performed for humane handling, caring for, and treating research animals (Protocol number MD-22-17143-5). Mice were kept in disposable cages supplemented with enrichment, and regular sterile food, water, and air were supplied separately in each cage. All cages were placed in specific pathogen-free (SPF) conditions during the experiment with controlled environmental conditions. The mice were left to acclimate for 3 days, after which treatment groups were randomized, and the littermates were evenly distributed in cages. Identification markings and shaving on dorsal flanks of already weighed mice were performed, and mice were infected. Following infection, twice daily, mice were given wet food and monitored based on parameters like body weight, activity level, fur, and eye appearance. As per the guidelines of the Institutional Animal Care Units of the Hebrew University’s School of Medicine, based on the above parameters, a scoring method was implemented to decide humane endpoints where mice were euthanized according to ethically approved procedures.

### Sublethal murine of human GAS soft tissue infection

The murine model of human soft-tissue infection with a sub-lethal dose (5 x 10^7^ CFU) of GAS strains was performed to determine CFU counts in soft skin and spleen samples. GAS strains were cultured in THY medium at 37°C and grown to OD_600_ = 0.3–0.4. Bacteria were then washed twice with sterile PBS and brought to a concentration of 5 x 10^7^ CFU in 100 μl, which were injected SC into the rear flank of mice. The actual bacterial inoculum was confirmed by counting CFU on blood agar plates of serially diluted final suspension. At various time points, mice were euthanized by inhalation of isoflurane followed by cervical dislocation, and skin and spleen samples were collected. Skin tissue from the injection site was collected using a punch biopsy tool (Acu-Punch, Acuderm Inc.), and spleen samples were excised and transferred to 2 ml Eppendorf tubes containing 0.5 ml of sterile PBS. Tissues were homogenized, diluted, and plated on blood agar plates for all experiments, and CFUs were counted after overnight incubation at 37°C. CFU counts were normalized to the weight of the soft tissue. The CFU counts for S119 *gln*P^-^ and S119 *asn*B^-^ mutants were matched with parallel plating on erythromycin (1 µg/µl) supplemented THY agar plates to check the stability of insertional inactivation.

For determination of the lesion area, the dermonecrotic skin lesions were measured daily using a digital caliper (Bar Naor Ltd.). The lesion area was calculated with the formula A = (π/2)(length)(width) (Anand et al., 2021).

### Statistical analysis

GraphPad Prism version 10 software was used to plot results and perform statistical analysis. All values were represented as means ± standard deviation (SD). Data in bar graphs were analyzed using parametric unpaired two-tailed t-tests unless specified, and two-way ANOVA with Tukey post-tests and Mann– Whitney U test, where indicated. In all figures, P values were calculated to confirm the significance: *p < 0.05; **p < 0.01; ***p < 0.001; ****p < 0.0001.

---

**Table S1.** List of bacterial strains used in the study.

| STRAIN | DESCRIPTION | SOURCE |
| --- | --- | --- |
| *Streptococcal* Strains |  |  |
| S119 | An invasive strain isolated from human blood in 2008 |  |
| S119 $\Delta covS$ | | This study |
| S119 $covR$ | | This study |
| S119 $\Delta asnA$ | | This study |
| S119 $\Delta rocA$ | | This study |
| S119 $\Delta asnA$ - pLZ *asnA* | | This study |
| S119 $\Delta asnA$ *R101K | | This study |
| S119 $\Delta asnA$ - pLZ *asnA* *R100K | | This study |
| S119 $glnP^-$ | | This study |
| S119 $glnP^-$ - pLZ *glnP* | | This study |
| S119 $asnB^-$ | | This study |
| S119 $asnB^-$ - pLZ *asnB* | | This study |
| GAS 854 | M1T1 GAS clinical isolate from a patient with a retroperitoneal abscess | <sup>4</sup> |
| GAS 854 *asnA*<sup>-</sup> |  | This study |
| GAS 5448 | M1T1 GAS clinical isolate from a patient with necrotizing fasciitis and toxic shock syndrome | <sup>5</sup> |

**Table S2.**
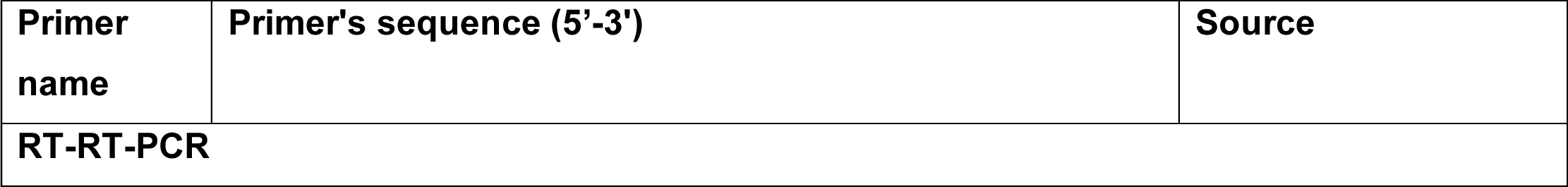
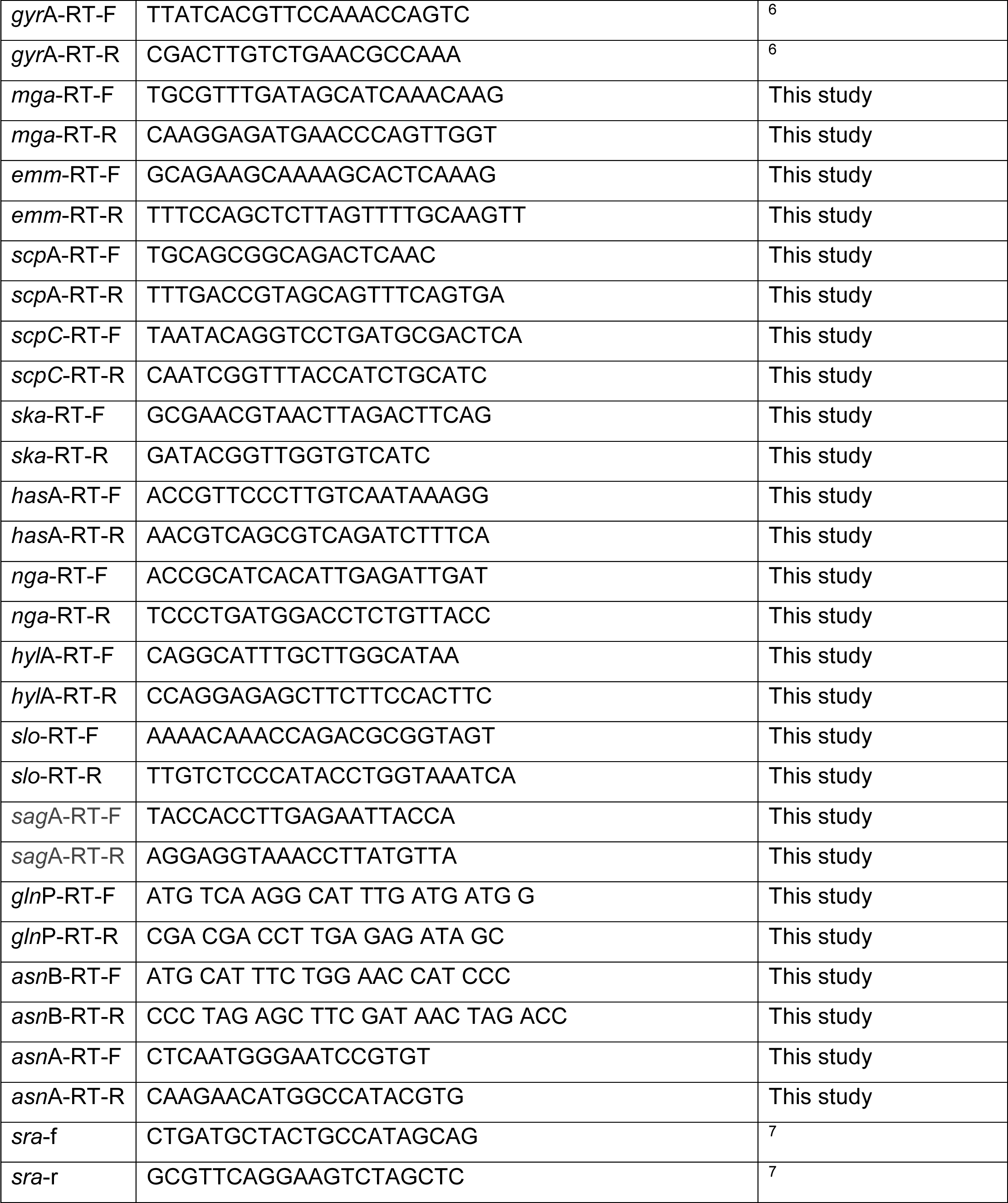
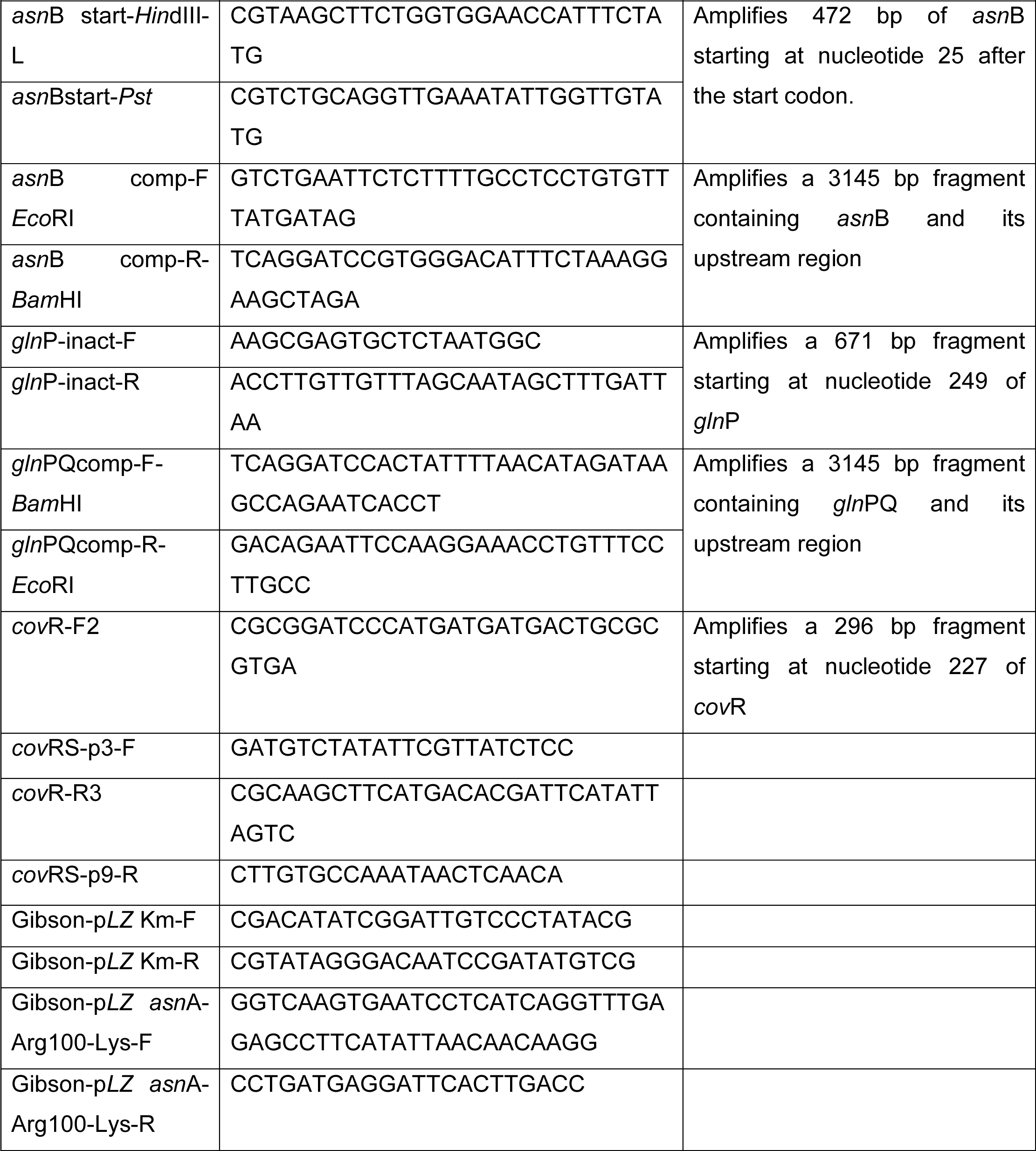
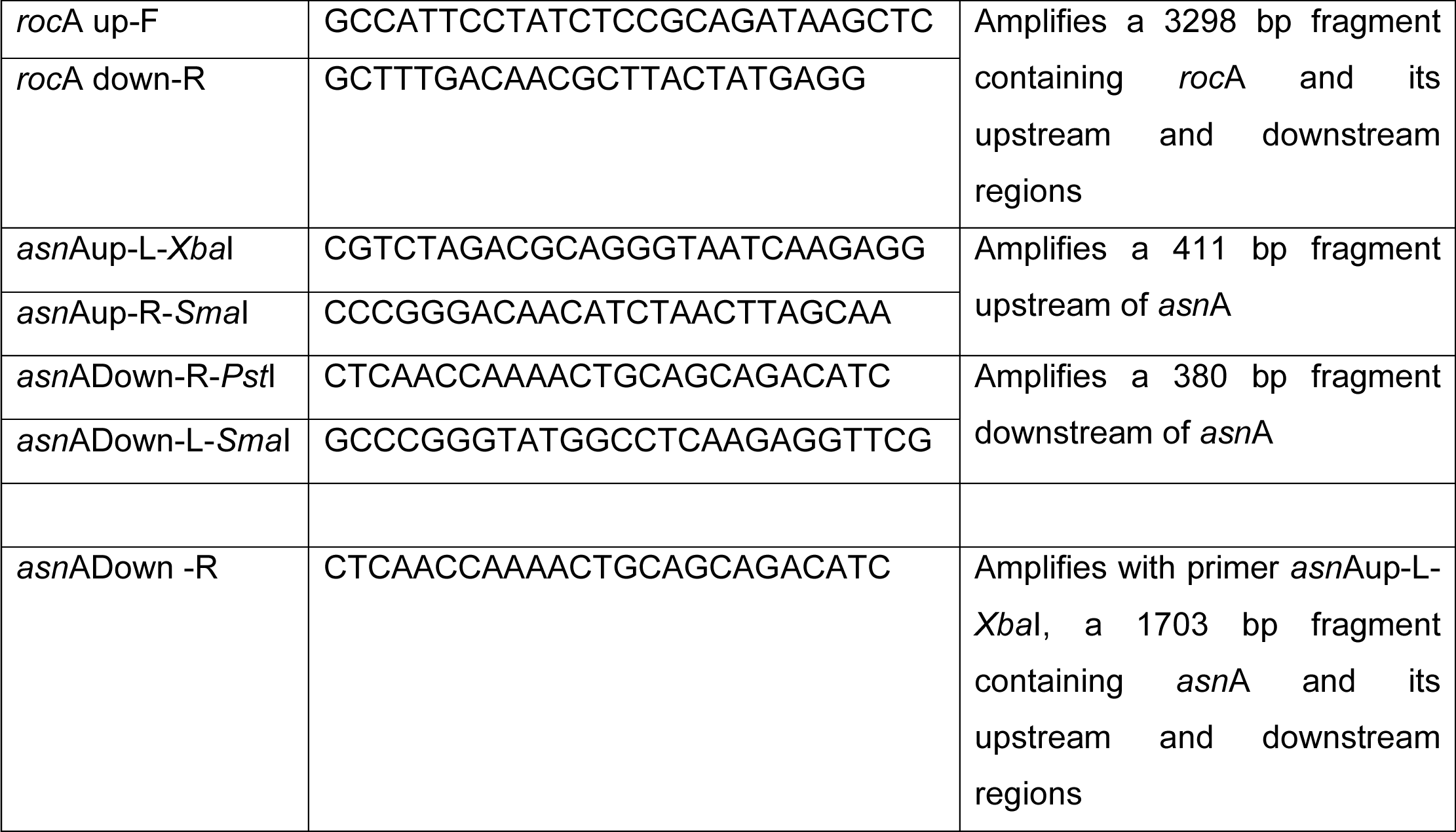
List of qRT-PCR primers used in the study.

**Table S3.**
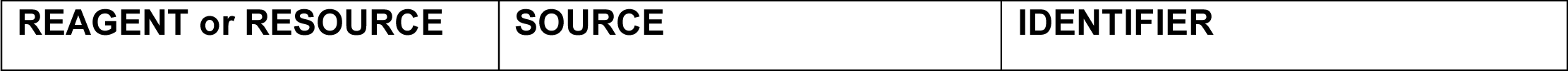
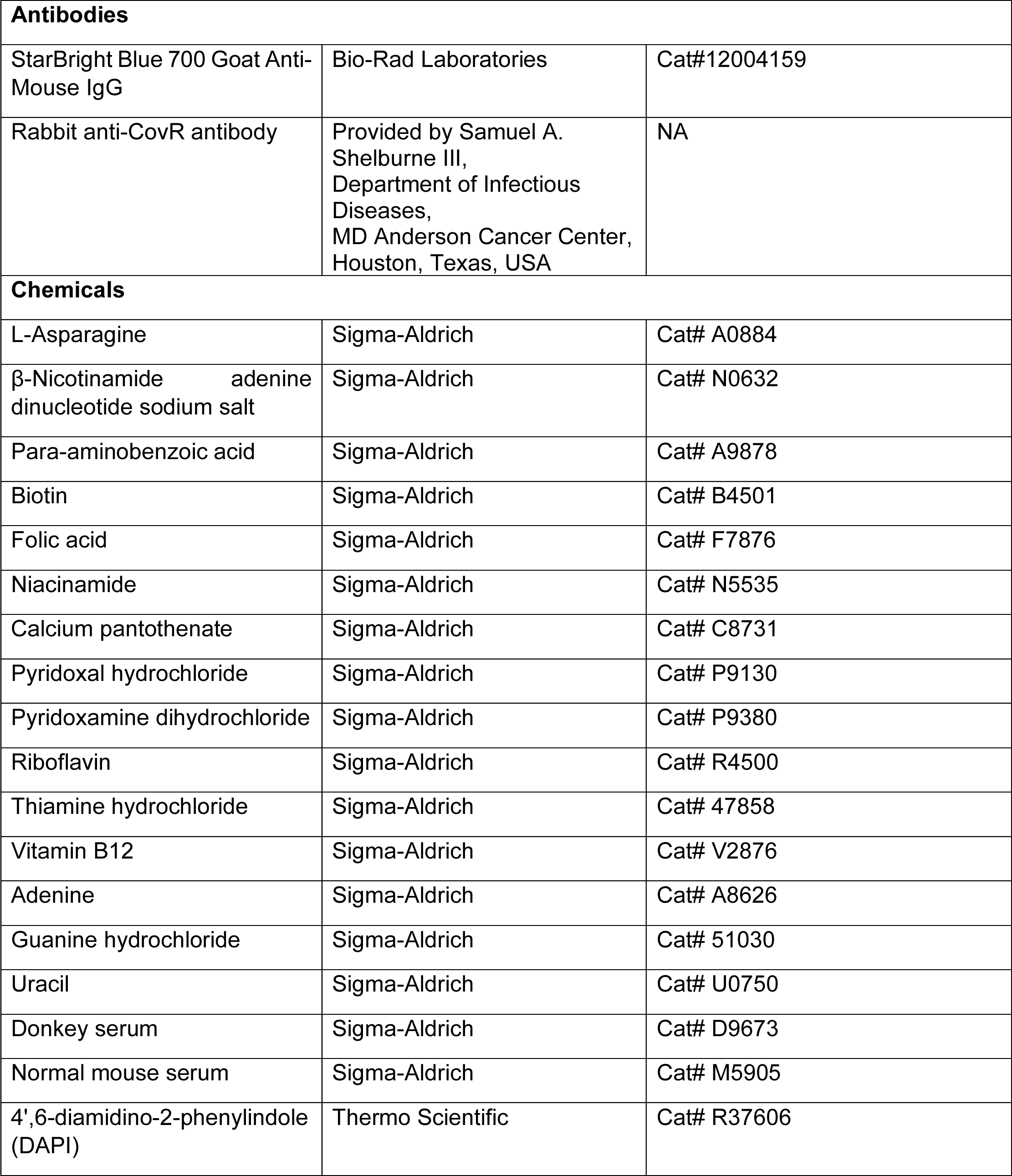
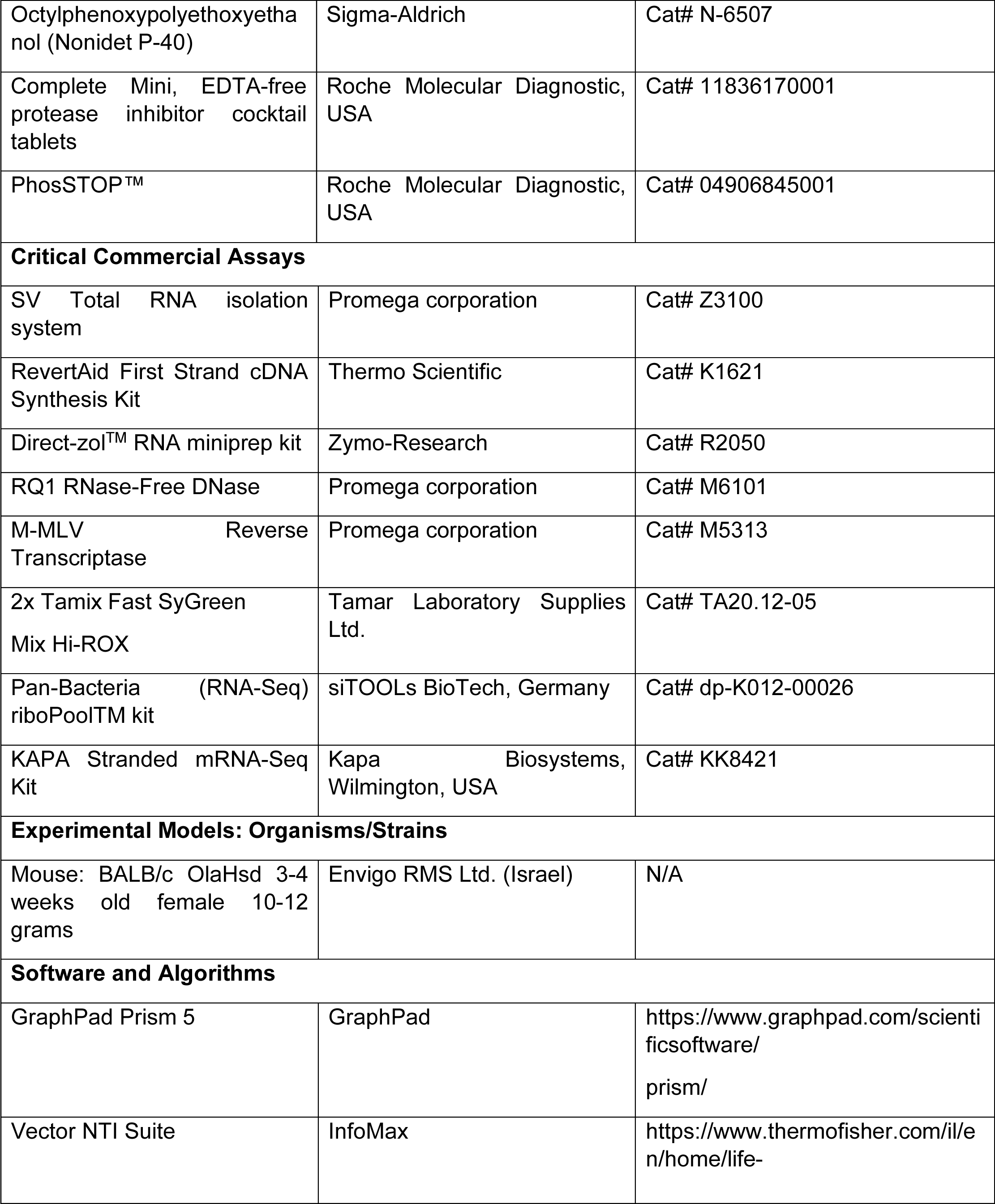
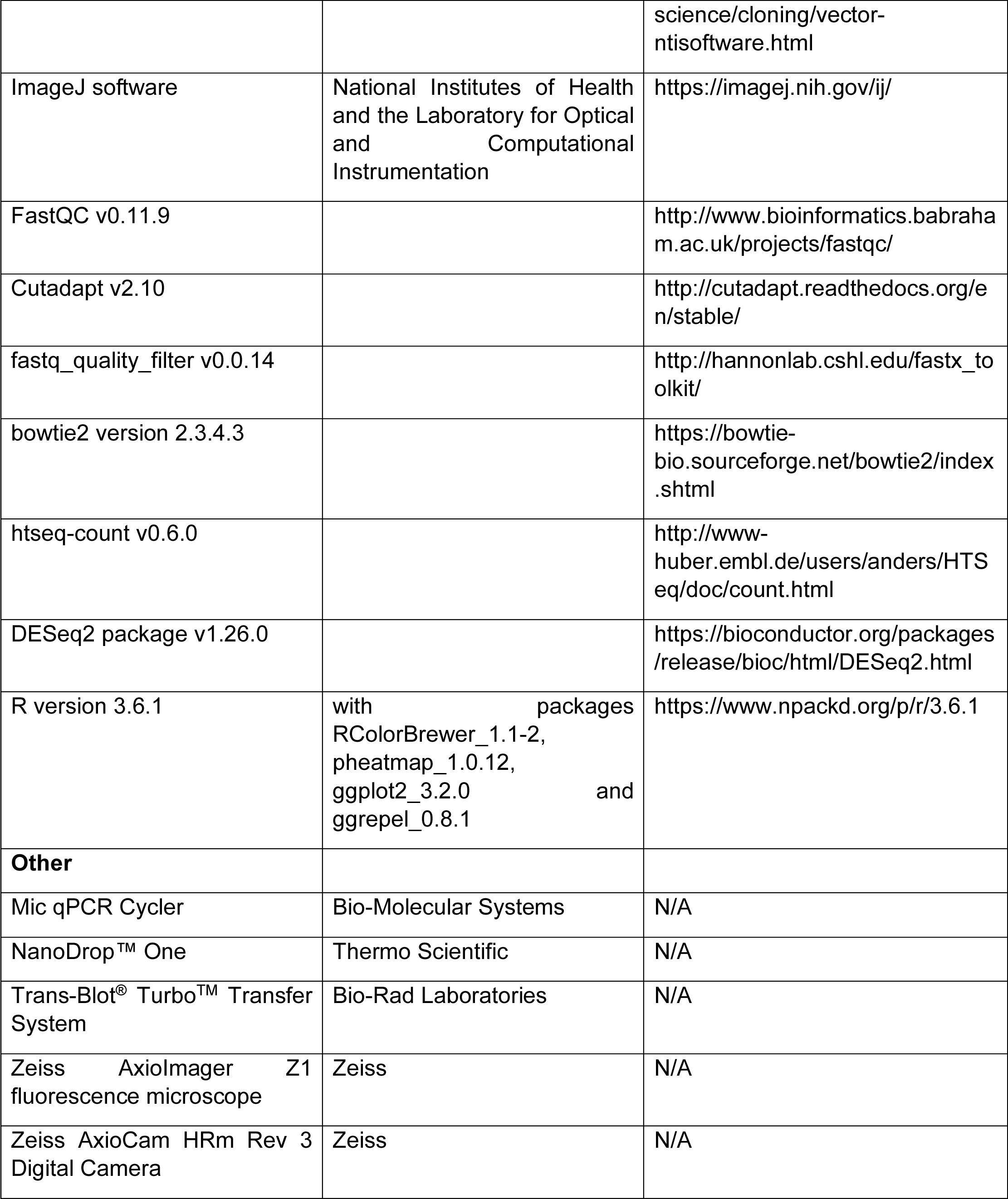
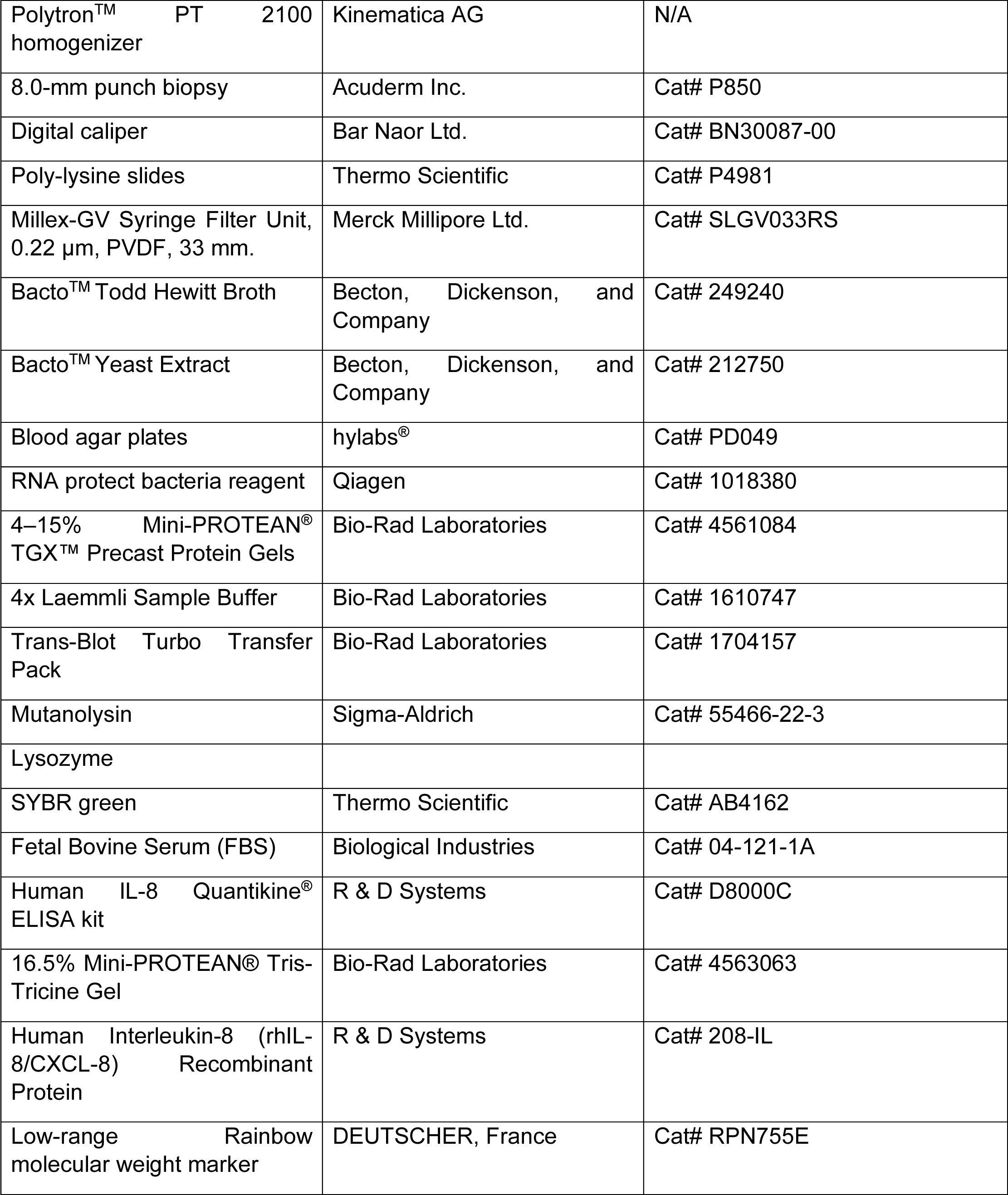
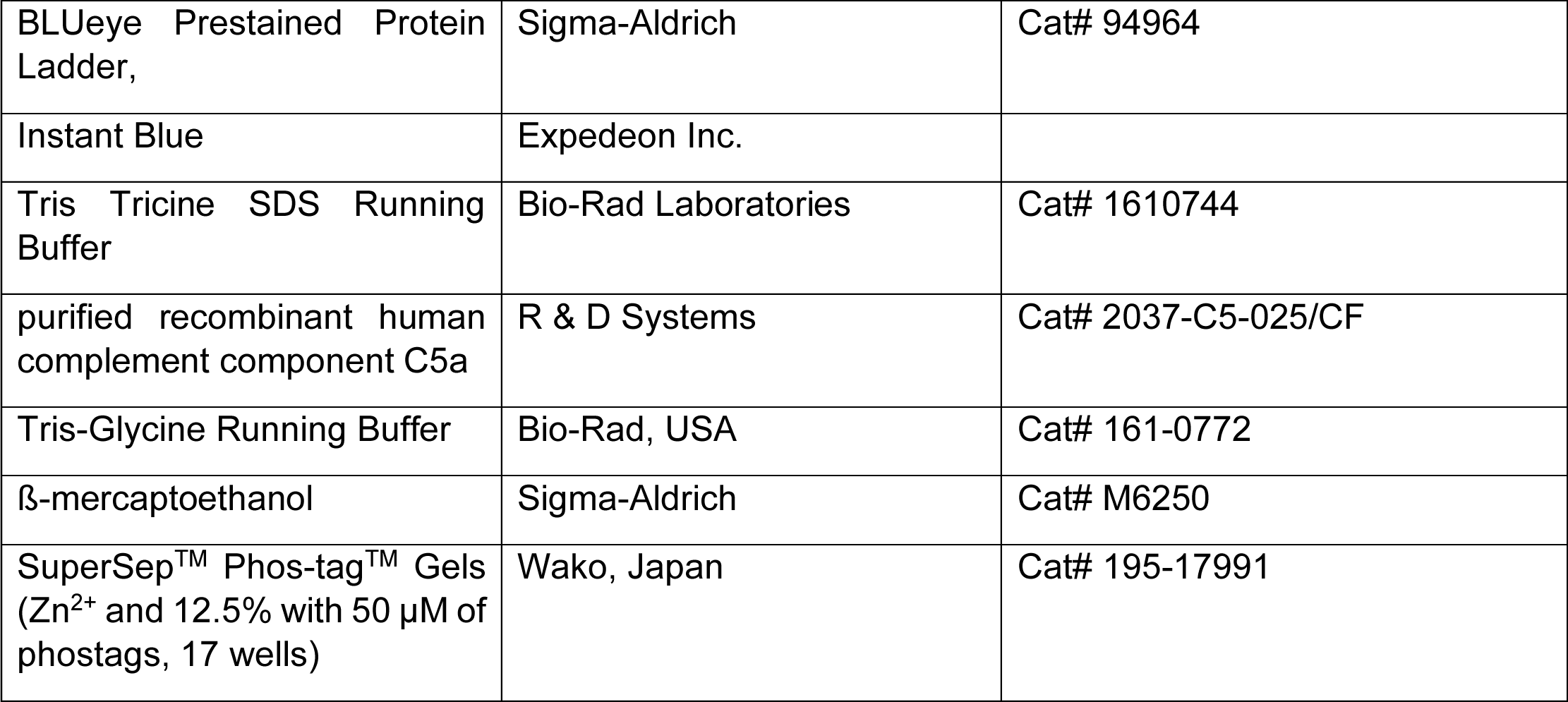
List of resources/reagents used in the study.

## References

1. Barnett, T.C., Bowen, A.C. & Carapetis, J.R. The fall and rise of Group A Streptococcus diseases. Epidemiol Infect 147, e4 (2018).

2. Olsen, R.J. & Musser, J.M. Molecular pathogenesis of necrotizing fasciitis. Annu Rev Pathol 5, 1–31 (2010).

3. Vega, L.A., Malke, H. & McIver, K.S. Virulence-Related Transcriptional Regulators of Streptococcus pyogenes, in Streptococcus pyogenes: Basic Biology to Clinical Manifestations, Edn. 2nd. (eds. J.J. Ferretti, D.L. Stevens & V.A. Fischetti) (Oklahoma City (OK); 2022).

4. Walker, M.J. et al. Disease manifestations and pathogenic mechanisms of Group A Streptococcus. Clin Microbiol Rev 27, 264–301 (2014).

5. Schiavolin, L., Deneubourg, G., Steinmetz, J., Smeesters, P.R. & Botteaux, A. Group A Streptococcus adaptation to diverse niches: lessons from transcriptomic studies. Crit Rev Microbiol 50, 241–265 (2024).

6. Krall, A.S. et al. Asparagine couples mitochondrial respiration to ATF4 activity and tumor growth. Cell Metab 33, 1013–1026 e1016 (2021).

7. Ralph, A.P. & Carapetis, J.R. Group a streptococcal diseases and their global burden. Curr Top Microbiol Immunol 368, 1–27 (2013).

8. Brouwer, S. et al. Pathogenesis, epidemiology and control of Group A Streptococcus infection. Nat Rev Microbiol 21, 431–447 (2023).

9. Cole, J.N., Barnett, T.C., Nizet, V. & Walker, M.J. Molecular insight into invasive group A streptococcal disease. Nat Rev Microbiol 9, 724–736 (2011).

10. Martin, W.J. et al. Post-infectious group A streptococcal autoimmune syndromes and the heart. Autoimmun Rev 14, 710–725 (2015).

11. Cunningham, M.W. Molecular Mimicry, Autoimmunity, and Infection: The Cross-Reactive Antigens of Group A Streptococci and their Sequelae. Microbiol Spectr 7 (2019).

12. Bagcchi, S. Surge of invasive Group A streptococcus disease. Lancet Infect Dis 23, 284 (2023).

13. Freiberg, J.A. & Wright, P.W. What’s Hot This Year in ID Clinical Science. Clin Infect Dis (2024).

14. Kim, T.H. Toxic Shock Syndrome (TSS) Caused by Group A Streptococcus: Novel Insights Within the Context of a Familiar Clinical Syndrome. J Korean Med Sci 39, e154 (2024).

15. Nygaard, U. et al. Invasive group A streptococcal infections in children and adolescents in Denmark during 2022-23 compared with 2016-17 to 2021-22: a nationwide, multicentre, population-based cohort study. Lancet Child Adolesc Health 8, 112–121 (2024).

16. Brown, S.A., Palmer, K.L. & Whiteley, M. Revisiting the host as a growth medium. Nat Rev Microbiol 6, 657–666 (2008).

17. Baruch, M. et al. An extracellular bacterial pathogen modulates host metabolism to regulate its own sensing and proliferation. Cell 156, 97–108 (2014).

18. Anand, A. et al. Unfolded protein response inhibitors cure group A streptococcal necrotizing fasciitis by modulating host asparagine. Sci Transl Med 13 (2021).

19. Gjymishka, A., Su, N. & Kilberg, M.S. Transcriptional induction of the human asparagine synthetase gene during the unfolded protein response does not require the ATF6 and IRE1/XBP1 arms of the pathway. Biochem J 417, 695–703 (2009).

20. Graham, M.R. et al. Virulence control in group A Streptococcus by a two-component gene regulatory system: global expression profiling and in vivo infection modeling. Proc Natl Acad Sci U S A 99, 13855–13860 (2002).

21. Horstmann, N. et al. Phosphatase activity of the control of virulence sensor kinase CovS is critical for the pathogenesis of group A streptococcus. PLoS Pathog 14, e1007354 (2018).

22. Velarde, J.J., Ashbaugh, M. & Wessels, M.R. The human antimicrobial peptide LL-37 binds directly to CsrS, a sensor histidine kinase of group A Streptococcus, to activate expression of virulence factors. J Biol Chem 289, 36315–36324 (2014).

23. Rosinski-Chupin, I., Sauvage, E., Fouet, A., Poyart, C. & Glaser, P. Conserved and specific features of Streptococcus pyogenes and Streptococcus agalactiae transcriptional landscapes. BMC Genomics 20, 236 (2019).

24. Plainvert, C. et al. A Novel CovS Variant Harbored by a Colonization Strain Reduces Streptococcus pyogenes Virulence. J Bacteriol 205, e0003923 (2023).

25. van de Rijn, I. & Kessler, R.E. Growth characteristics of group A streptococci in a new chemically defined medium. Infect Immun 27, 444–448 (1980).

26. Gogos, A., Jimenez, J.C., Chang, J.C., Wilkening, R.V. & Federle, M.J. A Quorum Sensing-Regulated Protein Binds Cell Wall Components and Enhances Lysozyme Resistance in Streptococcus pyogenes. J Bacteriol 200 (2018).

27. Mikkat, S., Kreutzer, M. & Patenge, N. Dynamic Protein Phosphorylation in Streptococcus pyogenes during Growth, Stationary Phase, and Starvation. Microorganisms 12 (2024).

28. Finn, M.B., Ramsey, K.M., Dove, S.L. & Wessels, M.R. Identification of Group A Streptococcus Genes Directly Regulated by CsrRS and Novel Intermediate Regulators. mBio 12, e0164221 (2021).

29. Horstmann, N., Myers, K.S., Tran, C.N., Flores, A.R. & Shelburne Iii, S.A. CovS inactivation reduces CovR promoter binding at diverse virulence factor encoding genes in group A Streptococcus. PLoS Pathog 18, e1010341 (2022).

30. Norton, S.J. & Chen, Y.T. Beta-aspartylhydroxamic acid: its action as a feedback inhibitor and a repressor of asparagine synthetase in Lactobacillus arabinosus. Arch Biochem Biophys 129, 560–566 (1969).

31. Federle, M.J., McIver, K.S. & Scott, J.R. A response regulator that represses transcription of several virulence operons in the group A streptococcus. J Bacteriol 181, 3649–3657 (1999).

32. Trevino, J. et al. CovS simultaneously activates and inhibits the CovR-mediated repression of distinct subsets of group A Streptococcus virulence factor-encoding genes. Infect Immun 77, 3141–3149 (2009).

33. Fulyani, F., Schuurman-Wolters, G.K., Slotboom, D.J. & Poolman, B. Relative Rates of Amino Acid Import via the ABC Transporter GlnPQ Determine the Growth Performance of Lactococcus lactis. J Bacteriol 198, 477–485 (2016).

34. Fulyani, F. et al. Functional diversity of tandem substrate-binding domains in ABC transporters from pathogenic bacteria. Structure 21, 1879–1888 (2013).

35. Podbielski, A. & Leonard, B.A. The group A streptococcal dipeptide permease (Dpp) is involved in the uptake of essential amino acids and affects the expression of cysteine protease. Mol Microbiol 28, 1323–1334 (1998).

36. Mackay, G.M., Zheng, L., van den Broek, N.J. & Gottlieb, E. Analysis of Cell Metabolism Using LC-MS and Isotope Tracers. Methods Enzymol 561, 171–196 (2015).

37. Lubkowski, J. & Wlodawer, A. Structural and biochemical properties of L-asparaginase. FEBS J 288, 4183–4209 (2021).

38. Farahat, M.G., Amr, D. & Galal, A. Molecular cloning, structural modeling and characterization of a novel glutaminase-free L-asparaginase from Cobetia amphilecti AMI6. Int J Biol Macromol 143, 685–695 (2020).

39. Edwards, R.J. et al. Specific C-terminal cleavage and inactivation of interleukin-8 by invasive disease isolates of Streptococcus pyogenes. J Infect Dis 192, 783–790 (2005).

40. Lynskey, N.N. et al. Multi-functional mechanisms of immune evasion by the streptococcal complement inhibitor C5a peptidase. PLoS Pathog 13, e1006493 (2017).

41. Hidalgo-Grass, C. et al. A streptococcal protease that degrades CXC chemokines and impairs bacterial clearance from infected tissues. EMBO J 25, 4628–4637 (2006).

42. Horstmann, N. et al. Dual-site phosphorylation of the control of virulence regulator impacts group a streptococcal global gene expression and pathogenesis. PLoS Pathog 10, e1004088 (2014).

43. Horstmann, N. et al. Characterization of the effect of the histidine kinase CovS on response regulator phosphorylation in group A Streptococcus. Infect Immun 83, 1068–1077 (2015).

44. Ravins, M. et al. Murine Soft Tissue Infection Model to Study Group A Streptococcus (GAS) Pathogenesis in Necrotizing Fasciitis. Methods Mol Biol 2427, 185–200 (2022).

45. Pancholi, V. & Caparon, M. Streptococcus pyogenes Metabolism, in Streptococcus pyogenes: Basic Biology to Clinical Manifestations, Edn. 2nd. (eds. J.J. Ferretti, D.L. Stevens & V.A. Fischetti) (Oklahoma City (OK); 2022).

46. Khara, P., Mohapatra, S.S. & Biswas, I. Role of CovR phosphorylation in gene transcription in Streptococcus mutans. Microbiology (Reading) 164, 704–715 (2018).

47. Martinez-Reyes, I. et al. Mitochondrial ubiquinol oxidation is necessary for tumour growth. Nature 585, 288–292 (2020).

48. Cole, J.N. et al. M protein and hyaluronic acid capsule are essential for in vivo selection of covRS mutations characteristic of invasive serotype M1T1 group A Streptococcus. mBio 1 (2010).

49. Allen, U. & Moore, D. Invasive group A streptococcal disease: Management and chemoprophylaxis. Paediatr Child Health 15, 295–302 (2010).

50. Anaya, D.A. & Dellinger, E.P. Necrotizing soft-tissue infection: diagnosis and management. Clin Infect Dis 44, 705–710 (2007).

51. Dale, J.B. & Walker, M.J. Update on group A streptococcal vaccine development. Curr Opin Infect Dis 33, 244–250 (2020).

## References for materials and methods

1. van de Rijn, I. & Kessler, R.E. Growth characteristics of group A streptococci in a new chemically defined medium. Infect Immun 27, 444–448 (1980).

2. Horstmann, N. et al. Distinct single amino acid replacements in the control of virulence regulator protein differentially impact streptococcal pathogenesis. PLoS Pathog 7, e1002311 (2011).

3. Gao, J., Gusa, A.A., Scott, J.R. & Churchward, G. Binding of the global response regulator protein CovR to the sag promoter of Streptococcus pyogenes reveals a new mode of CovR-DNA interaction. J Biol Chem 280, 38948–38956 (2005).

4. Tran-Winkler, H.J., Love, J.F., Gryllos, I. & Wessels, M.R. Signal transduction through CsrRS confers an invasive phenotype in group A Streptococcus. PLoS Pathog 7, e1002361 (2011).

5. Zinkernagel, A.S. et al. The IL-8 protease SpyCEP/ScpC of group A Streptococcus promotes resistance to neutrophil killing. Cell Host Microbe 4, 170–178 (2008).

6. Eran, Y. et al. Transcriptional regulation of the sil locus by the SilCR signalling peptide and its implications on group A streptococcus virulence. Mol Microbiol 63, 1209–1222 (2007).

7. Belotserkovsky, I. et al. Functional analysis of the quorum-sensing streptococcal invasion locus (sil). PLoS Pathog 5, e1000651 (2009).

